# Simple Assay, Kinetics, and Biochemical Trends for Soil Microbial Catalases

**DOI:** 10.1101/2020.06.11.147595

**Authors:** Michael Chabot, Ernesto Morales, Jacob Cummings, Nicholas Rios, Scott Giatpaiboon, Rakesh Mogul

## Abstract

In this report, we expand upon the enzymology and biochemical ecology of soil catalases through development and application of a simple kinetic model and assay based upon volume displacement. Through this approach, we (A) directly relate apparent Michaelis-Menten terms to the catalase reaction mechanism, (B) obtain upper estimates of the intrinsic rate constants for the catalase community 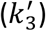 and moles of catalase per 16S rRNA gene copy number, (C) utilize catalase specific activities (SAs) to obtain biomass estimates of soil and permafrost communities (LOD, ~10^4^ copy number gdw^−1^), and (D) relate kinetic trends to changes in bacterial community structure. This model represents a novel approach to the kinetic treatment of soil catalases, while simultaneously incorporating barometric adjustments to afford comparisons across field measurements. As per our model, and when compared to garden soils, biological soil crusts exhibited ~2-fold lower values for 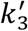, ≥10^5^-fold higher catalase moles per biomass (250-1200 zmol copy number^−1^), and ~10^4^-fold higher SAs per biomass (74-230 fkat copy number^−1^). However, the highest SAs were obtained from permafrost and high-elevation soil communities (5900-6700 fkat copy number^−1^). In sum, these total trends suggest that microbial communities which experience higher degrees of native oxidative stress possess higher basal intracellular catalase concentrations and SAs per biomass, and that differing kinetic profiles across catalase communities are indicative of phylum and/or genus-level changes in community structure. For microbial ecology, therefore, these measures effectively serve as markers for microbial activity and abundance, and additionally provide insights into the community responses to exogenous stress.

**Importance:** The efficient management of oxidative stresses arising from environmental pressures are central to the homeostasis of soil microbial communities. Among the enzymes that manage oxidative stress are catalases, which degrade hydrogen peroxide into oxygen gas and water. In this report, we detail the development and application of a simple kinetic model and assay to measure catalase reaction rates and estimate soil biomass. Our assay is based upon volume displacement, and is low-cost, field-amenable, and suitable for scientists and educators from all disciplines. Our results suggest that microbial communities that experience higher degrees of native oxidative stress possess higher basal intracellular catalase concentrations and specific activities when expressed per biomass. For microbial ecology, therefore, these measures serve as biochemical markers for microbial activity and abundance, and provide insights into the community responses to exogenous stress; thereby providing a novel means to study active microbial communities in soils and permafrost.

## Introduction

The efficient management of oxidative stresses arising from reactive oxygen species are central to homeostasis (1–3). For soil microbial communities, reactive oxygen species arise from exposures to ultraviolet radiation, oxidants, and desiccation, as well as aerobic respiration and photosynthesis (4–6). Among the array of intracellular enzymes that manage oxidative stress are the catalase enzymes, which degrade hydrogen peroxide (H_2_O_2_) into oxygen gas and water (7–9). The activities of catalase enzymes are linked with all taxonomic domains, with catalase genes additionally being present in facultative and obligate anaerobic microorganisms (10–12).

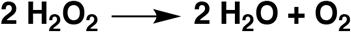

Catalases are organized into three major classes comprising monofunctional enzymes containing either iron-heme (Fe(heme)) or binuclear manganese (Mn_2_) metal cofactors (EC 1.11.1.6), and bifunctional catalase-peroxidases containing Fe(heme) cofactors (EC 1.11.1.21). As diagramed below, each enzyme catalyzes the disproportionation of H_2_O_2_ in two broad reaction steps, which commences with efficient capture of H_2_O_2_ to yield oxidized metal cofactors (13, 14), and concludes with liberation of oxygen with reaction rates nearing the limits of diffusion (~10^7^ M^−1^ s^−1^ for soluble catalases) (15, 16).

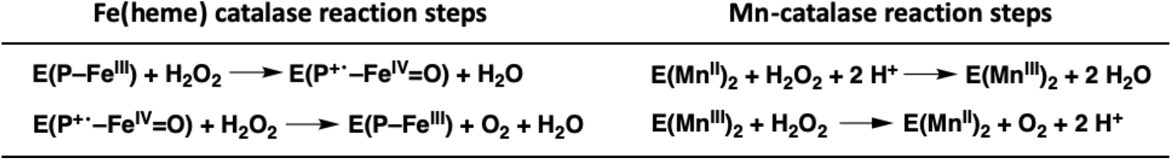

For the Fe(heme) catalases, capture of H_2_O_2_ results in irreversible formation of an oxyferryl porphyrin (P) radical cation, referred to as Compound I (E(P^+•^–Fe^IV^=O)), and liberation of water (17–19). For the Mn catalases (when beginning in the (Mn^II^)_2_ state), capture of H_2_O_2_ yields an oxidized manganese complex ((Mn^III^)_2_), and irreversible liberation of two waters (11, 12, 20). Productive completion for both mechanisms concludes with acquisition of a second H_2_O_2_, reduction of the metal centers, and liberation of O_2_ (and an additional water molecule for the Fe(heme) catalases). Potential non-productive catalytic steps include formation of a potential hydroxyferryl species (Compound II), which ultimately reverts back to the active enzyme (18).

For microbial ecology, measures of these reactions can serve as biochemical markers for intracellular biological activity, and provide insights into the community responses to exogenous stress. Among the more commonly used measures for catalase reactions include potassium permanganate (KMnO_4_) titrations (21, 22) and enzyme-linked colorimetric assays (23–25). Through straightforward, but relatively lengthy procedures, these methods provide single-time point rates, which are then converted to specific activities using soil mass. For KMnO_4_ titrations, however, the resulting specific activities are not necessarily amenable to units of enzyme activity (*e.g.*, Units or katals), as side reactions with soil organics prevent direct stoichiometric relationships. Moreover, titrations with KMnO_4_ introduce several chemical safety issues including proper storage of solutions and chemical waste during field campaigns, including waste disposal.

Alternatively, catalase rates can be measured via manometric and electrochemical approaches, which are amenable to kinetic measures of product formation (O_2_) – or multiple measures of reaction progress over time. Manometric approaches are not well represented in the current literature, with most (if not all) reported analytical assemblies being ill-suited for field work or field campaigns (26–28). In contrast, electrochemical and gasometric measures for purified catalases, biological extracts, and finely sieved plant-based powders are fairly well described (29–31); however, reports detailing the analyses of soil-based samples are rather limited (31, 32).

Hence, in this report, we expand upon the enzymology and biochemical ecology of catalases through development and application of a simple kinetic model and assay based upon volume displacement. This model represents a novel approach to the kinetic treatment of soil catalases, and explicitly correlates to the enzyme reaction mechanism for each type of catalase. Moreover, our kinetic assay is low-cost, rapid, field-amenable, applicable to a variety of environmental catalases, and suitable for scientists and educators from all disciplines. In addition, by incorporating biomass and barometric terms, our model affords upper estimates of the intrinsic rate constants and moles of catalase per soil mass and biomass, while allowing for the comparison of measurements obtained across geographies. In summary, our results suggest that microbial communities from biological soil crusts, high elevation soils, and permafrost experience substantial degrees of native oxidative stresses, and accordingly exhibit high catalase activities and abundances per 16S rRNA gene copy number.

## Materials & Methods

### Materials

Hydrogen peroxide was purchased as 30% *w/w* non-stabilized solutions (Sigma-Aldrich, St. Louis, MO) and 3% *w/w* stabilized solutions (CVS Pharmacy). Non-stabilized hydrogen peroxide, which did not contain chemical stabilizers, was immediately stored as aliquots at – 20°C. Additional reagents included bovine liver catalase (Sigma-Aldrich), HEPES (4-(2-hydroxyethyl)-1-piperazineethanesulfonic acid) (VWR, Radnor, PA), NaCl (VWR), phosphate-buffered saline (PBS) tablets (VWR), and 10x PBS solution (100 mM potassium phosphate, 100 mM NaCl, pH 7.4) (VWR). All solutions were prepared in ultrapure water (18 MΩ cm^−1^), and sterile filtered (0.22 μm syringe filters) or autoclaved.

The volume displacement (VD) apparatus (**Diagram 1 & Figures S1A-C**) was assembled using common laboratory supplies; no major purchases were required for assembly or reliable long-term operation of the apparatus. Materials included two 50 mL conical tubes, a tube rack, <3 ft of Tygon tubing (1/8 in, 1/4 in, 1/16 in), 1 one-hole rubber stopper (1.4 × 1.1 in), 1 two-hole rubber stopper (1.4 × 1.1 in), parafilm (optional), a 15 mL graduated cylinder (or a 15 or 50 mL conical tube), a mass balance (minimum accuracy of 0.01 g), a stir plate, 3 mm magnetic stir bar, and a stopwatch.

The electrochemical (EC) apparatus (**Figure S1D**) was assembled using a O_2_ Gas Sensor (O2-BTA) and LabQuest 2 (LABQ2) data logger from Vernier Software & Technology (respective list costs of $199 and $329). Additional materials included a support stand (~15-24 in tall), 3-prong clamp, magnetic stir plate, 3 mm stir bar, 50 mL conical tube (mixing chamber), one and two-hole rubber stoppers, Tygon tubing, two plastic stopcocks, and parafilm.

### Soil Samples

Soil samples, as summarized in **Table S1**, were obtained from ecoregions spanning differing hemispheres, elevations, annual rainfalls, and irrigation frequencies. Samples were collected in duplicate or triplicate using sterile or cleaned spatula, stored in sterile 15 or 50 mL conical tubes, and analyzed immediately using field devices, within 1-5 days in makeshift or field station laboratories, or within 2 weeks in a formal laboratory.

Garden and landscaped soils from sites of differing irrigation frequencies were obtained on the campus of Cal Poly Pomona (CPP) in Pomona, CA, USA between September 2017 and January 2018. Samples collected from (or near) the BioTrek Ethnobotany Garden (251 m; 34.057220, −117.826476) included (A) bare damp topsoils which were irrigated 2-3 times/week and adjacent to extensive plant coverage (CPP.BioTrek.Garden), (B) damp soils covered by substantial leaf litter which were irrigated 2-3 times/week and adjacent to extensive plant coverage (CPP.BioTrek.LeafLitter), and (C) non-landscaped soils that were fully removed from surrounding plant cover and subjected to no regular irrigation (CPP.BioTrek.DryPatch). Additional samples from CPP included bare dry topsoils that were irrigated ≤1/week, and adjacent to Japanese Iris (CPP.EnvDes.DryPatch; 255 m, 34.057205, −117.827231) and an Oak tree (CPP.Quad.DryPatch; 234 m; 34.058643, −117.823691). Averaged dry weights were calculated in triplicate after heating samples at 100 °C for 48 h (VWR 1530 Incubator).

Cold, dry, and high elevation soils were obtained from the Tibetan Plateau in Ladakh, India (3300-5400 m) in August 2017 (33). Samples were collected between sparse and small plant coverage near Tso Kar Lake (4592 and 4594 m; 33.315731, 77.955639), Taglang La (5383 m; 33.508517, 77.771442), and Khardung La (5359 m; 34.279661, 77.603806). Samples were also collected from a regularly irrigated vegetable garden at the Silk Road Cottages in Sumur (3300 m; 34.624553, 77.622154).

Permafrost samples were collected near Tso Kar Lake (5350 m; 33.315731, 77.955639) in Ladakh, India in August 2017, and analyzed within 1 day of acquisition. Alaskan permafrost samples (139 m; 64.951, −147.621) were collected in 2012 by Mackelprang *et al*. at the United States Army Cold Regions and Research and Engineering Laboratory (CRREL) permafrost tunnel in Fox, Alaska; samples were stored at −20 °C, and analyzed in 2016 for this study.

Black-crusted biological soil crusts (BSCs) samples were obtained in March 2016 from the Mojave National Preserve near Baker, CA, USA (off of Kelbaker Rd.). Samples were collected from areas of high (685 m; 35.198900, −115.870850) and intermediate (450 m; 35.255217, −115.957150) surface coverages (or surface densities) for the BSCs. Samples included the BSC topsoils (top 1 cm of the black crust) and BSC subsurfaces (the following 1 cm of crust).

Alkaline evaporate samples were obtained from Soda Lake (283 m; 35.148570, – 116.091136) in March 2016 from the Mojave National Preserve near Baker, CA, USA (off of Zzyzx Rd.). Samples from a ~30 cm depth profile were collected using a metal coring device. However, due to the loosely associated evaporates, coring resulted in significant compression of the sample; thereby, yielding a final ~10 cm core. Sub-samples were extracted at 1 cm increments, andaqueous extracts of the surface samples provided pH values of ≥11.

### Kinetic Assays with Soil Samples

Reaction rates were measured by volume displacement (VD) and electrochemical detection (EC). Devices for VD and EC were assembled as described in the **Supplementary Materials**. As per **Diagram 1** (and **Figures S1A-C**), the VD apparatus comprised sequentially connected chambers respectively used for mixing, water displacement, and water collection. For VD, the mass of collected water over time was proportional to the rate of product formation.

Kinetic analyses were performed and replicated on all described soils. For VD, this included BSCs studied in laboratory settings (n=3), BSCs studied in the field (n=2), dry high-elevation soils from Ladakh (n=3), CPP soils (n=3), Ladakh permafrost (n=2), Alaskan permafrost studied at 22 °C (n=9), and Alaskan permafrost studied at 4 °C (n=3). For EC (**Figure S1D**), this included BSCs studied in laboratory settings (n=5), BSCs studied in the field (n=2), Alaskan permafrost studied at 22°C (n=3), and Soda Lake samples (n=2). For all studies, a new conical tube was used for each reaction.

**Diagram 1.**
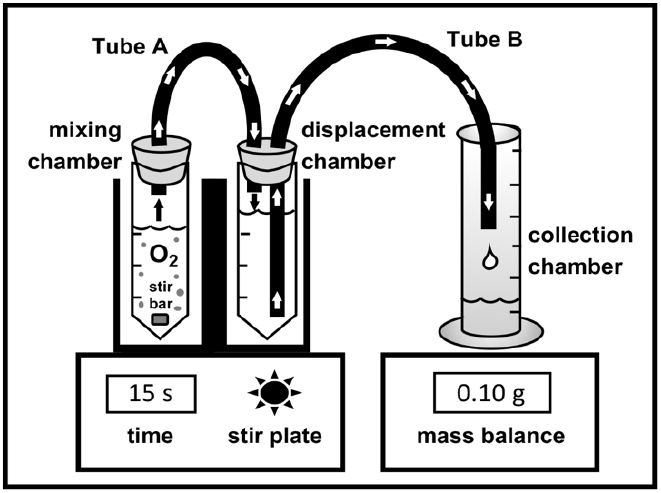
Volume displacement device with sequentially connected mixing, displacement, and collection chambers.

Samples were prepared by initially parsing the soils to remove rocks, leaves, and other debris. Most samples were subsequently crushed for 30 s using a mortar and pestle. Prepared soils (1-10 g) were transferred to the VD and EC mixing chambers using ~ 1.0 g for garden and landscaped soils, high-elevation dry soils, or BSCs, 1-2 g for Alaskan permafrost, and 10 g for Tso Kar permafrost. Soil samples were resuspended in 50 mM HEPES (pH 7.5) or 1x PBS, and vigorously mixed using a 3 mm stir bar and magnetic stir plate (medium setting). All reactions were 30 mL in final volume, and initiated by the addition of substrate using non-stabilized or stabilized formulations of hydrogen peroxide (H_2_O_2_).

For Michaelis-Menten studies, BSCs and garden soils were resuspended in solutions containing 3 mL 10x buffer solution (500 mM HEPES, pH 7.5) and sufficient ultrapure water to achieve a final reaction volume of 30 mL. Enzymatic reactions were initiated by the addition of 6.1 μL – 2.1 mL 30% *w/w* non-stabilized H_2_O_2_, which amounted to a range of final concentrations of 20-700 mM H_2_O_2_. For specific activity measurements, final H_2_O_2_ concentrations were 330 mM (or 1% *w/w*). When using non-stabilized H_2_O_2_, soil samples were resuspended in 29 mL buffer, and reactions initiated with 1 mL 30% *w/w* non-stabilized H_2_O_2_. When using stabilized H_2_O_2_, soil samples were resuspended in 20 mL buffer, and reactions initiated with 10 mL 3% *w/w* stabilized H_2_O_2_.

Upon addition of H_2_O_2_, the mixing chambers for VD and EC were rapidly sealed with the respective rubber stoppers. For VD, reactions were monitored by following the change in mass in the collection chamber every 15 s for at least 120 s. For EC, reactions were monitored by following the change in % O_2_ every 2 s for at least 300 s. Kinetic assays were conducted at 22 °C in a formal chemistry laboratory at CPP (BSCs, CPP soils, and Alaskan permafrost), at 4 °C using a reach-in refrigerator at CPP (Alaskan permafrost), at 21 °C in a makeshift laboratory in Ladakh, India (Ladakh dry soils and permafrost), and at 28-32 °C in the field station laboratory at the California State University (CSU) Desert Studies Center (BSCs and Soda Lake).

Reaction rates for VD were calculated by linear regression between 15-75 s (or linear portion of the rate plot) using a R^2^ value of ≥0.98 (Microsoft Excel). Rates were expressed as the grams of water displaced per second (g H_2_O displaced s^−1^). For EC experiments, regressions (LabQuest 2) were performed over the linear portion of the plots (minimum of 20 s or 10 data points), and rates expressed as the % O_2_ detected per minute (% O_2_ min^−1^). As described in below, all rates were converted to μmoles H_2_O_2_ consumed per second (μmoles s^−1^, μkat) and expressed as specific activities. For Michaelis-Menten analyses, substrate concentrations were normalized to soil mass (mM g^−1^), and non-linear least squares regressions (Microsoft Excel) provided the apparent (*) parameters of 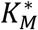 and 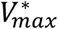.

### Field Assays

For field assays (**Figure S1C**), the VD and EC devices relied on a battery-powered stir plate (MagneStir) and analytical balance (Digital Scale, China). During an expedition in the Mojave National Preserve (March 2016), assays were conducted in the field on BSCs immediately after sampling. Supplies were transported to the sampling site using a medium size plastic container (~16 gal capacity), which alternatively served as a housing for the field apparatus during the analyses (**Figure S1E**). To ensure reagent integrity, 30% *w/w* non-stabilized H_2_O_2_ solutions were stored on ice packs in a foam cooler (while the 3% *w/w* stabilized H_2_O_2_ solutions were stored at ambient conditions). To minimize the impacts of wind (especially during weighing of samples), an umbrella was held in position near the VD and EC assemblies. At the CSU Desert Studies Center, electric-powered devices were used when conducting experiments in the field station laboratory (28-32 °C). During a field campaign in Ladakh, India (August 2017), VD assays were conducted in the city of Leh (3409 m) in the meeting/conference room at the Mogol Hostel (21 °C). For transport to India, supplies (*e.g.*, disassembled apparatus, pipets, and battery-operated devices) were safely organized in a travel suitcase/backpack and transported as checked-in baggage during international and domestic flights. Assays in Ladakh were conducted using 1x PBS (prepared using PBS tablets and bottled drinking water) and stabilized 3% *w/w* H_2_O_2_ (obtained from a pharmacy in Delhi, India); all solutions were stored at ambient conditions.

### Unit Conversions & Expression of Specific Activity

Reaction rates from VD and EC experiments were converted to SI units of μkatal (μkat), or the micromoles of substrate consumed per second (where 1 μkat = 60 Units), and expressed as specific activities by normalizing to grams of fresh soil sample (g^−1^), gram dry weight (gdw^−1^), 16S rRNA gene copy numbers (copy number^−1^), or grams of protein (mg^−1^). For VD experiments, measured reaction rates (g H_2_O displaced s^−1^) were firstly divided by the density of water to provide the volume of water displaced per second (mL H_2_O displaced s^−1^). Due to the proportional displacement of water by the evolved oxygen gas, the volume of displaced water by the reaction (per second) was assumed equal to the volume of oxygen released into the gas phase (per second). The volume of oxygen released (mL O_2_ released s^−1^) was then converted to the moles of oxygen released (*n*) through the ideal gas law (*PV* = *nRT*) using the gas constants of 8.314 L kPa K^−1^ mol^−1^ or 0.08205 L atm K^−1^ mol^−1^.

To enact the conversion to moles, the partial pressure of oxygen 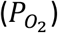 was obtained using Dalton’s law (**Equation 1**), which corrected for the impacts of water vapor 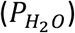 on the total pressure (*P*_*T*_). In turn, the partial pressure of water 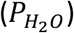 was obtained from **Equation 2**, where the saturation vapor pressure 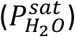 and relative humidity (*RH*) at the respective temperatures and sites of experimentation were obtained from online resources (*e.g.*, Weather Underground and Google Maps), CRC handbook, or hand-held devices.

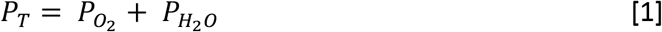

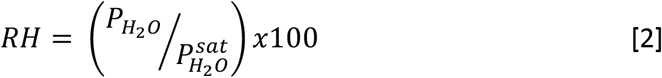

To account for the impacts of elevation (*z*; meters) and temperature (T; Kelvin), the total pressure was adjusted using the barometric formula outlined in **Equation 3** (34), where *P*_3_ was assigned as the station pressure (*P*_*z*_) and calculated by adjusting the sea level pressure (*P*_0_), or equivalent pressure at sea level (obtained from Weather Underground). Simplification to **Equation 4** was afforded by combining all constant terms in to yield 3.417×10^−2^ K m^−1^, which included the molar mass of Earth’s air (M; 28.97 g mol^−1^; assuming 78% N_2_, 21% O_2_, and 1% Ar), acceleration of gravity (g; 9.80665 m s^−2^), and gas constant (R; 8314 g m^2^ s^−2^ K^−1^ mol^−1^, or 8.314 JK^−1^ mol^−1^, where J = kg m^2^ s^−2^). Upon conversion using the ideal gas law, the moles of oxygen released per second (moles O_2_ released s^−1^) were then converted to the micromoles of H_2_O_2_ (or substrate) consumed per second (μkat; or μmoles H_2_O_2_ consumed s^−1^) using the reaction stoichiometry. All rates were expressed as specific activities (μkat g^−1^, μkat gdw^−1^, or μkat copy number^−1^).

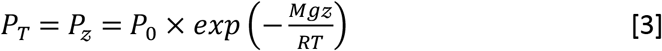

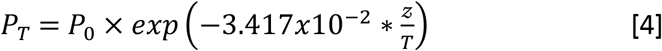

For EC experiments, measured rates (%O_2_ min^−1^) were re-expressed (after dividing by 100) as the ‘volume of oxygen’ measured (cm^3^) per ‘volume of the total headspace’ (cm^3^) per second (cm^3^ O_2_ cm^−3^ s^−1^). In turn, the rate was transformed to the ‘volume of oxygen’ measured per second (cm^3^ O_2_ s^−1^) by multiplying by the ‘volume of the total headspace’. For the EC apparatus, the estimated total headspace (76 cm^3^, 76 mL) encompassed the mixing chamber headspace (~20 mL), connective tubing (~2 mL), and internal sensor volume (~54 mL). Conversion of volume to the ‘moles of oxygen’ measured per second (moles O_2_ s^−1^) was thus afforded using the ideal gas law and 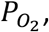, as described. All EC rates were converted to micromoles of H_2_O_2_ consumed, and expressed as specific activities (μkat g^−1^ or μkat mg^−1^).

### Kinetic Assays with Soluble Enzymes

Specific activities were measured for bovine liver catalase and clarified extracts of *Acinetobacter radioresistens* 50v1, a hydrogen peroxide-tolerant Gram-negative bacterium. For comparative purposes, the catalase specific activities were analyzed by VD, EC, and absorbance spectroscopy. Stock solutions of 10 mg/mL bovine liver catalase were prepared in 50 mM HEPES (pH 7.5). Extracts of *A. radioresistens* 50v1 were prepared and characterized as described (9). Reactions with soluble enzymes were conducted in 50 mM HEPES (pH 7.5) using 20 mM non-stabilized H_2_O_2_.

For VD and EC assays, reactions were 30 mL in final volume and contained 2.1 nM (0.50 μg/mL) bovine liver catalase, or 0.015 mg/mL (or 3.0 mL) of the clarified extract from *A. radioresistens* 50v1. Enzymatic reactions were initiated by addition of a final concentration of 20 mM H_2_O_2_. Reactions were monitored and rates calculated as described. All reactions rates were reproduced (bovine liver catalase, n=6 for VD and n=5 for EC; for the 50v1 extract, n=3 for VD and n=2 for EC).

For absorbance spectroscopy (Beckman DU-640), solutions were 1 mL in final volume, and contained 50 mM HEPES (pH 7.5) and 20 mM H_2_O_2_. Enzyme reactions were initiated by the addition of 2.1 nM (0.50 μg/mL) bovine liver catalase, or 0.015 mg/mL (or 100 μL) of the 50v1 extract (final concentrations). Reaction progress was monitored by following the change in absorbance every 2 s at 240 nm (for at least 60 s), where decreases in absorbance correlated to the consumption of H_2_O_2_ by catalase. Reaction rates were calculated using R^2^ values of ≥0.95 over a *minimum* of 14 s (or 7 data points); however, in practice, most regressions were R^2^ ≥0.99. Reaction rates (μmol s^−1^) were calculated using the molar extinction coefficient for H_2_O_2_ (43.6 M^−1^ cm^−1^), and total reaction volume (1 mL). Specific activities were expressed per mg of protein (μkat mg^−1^), and all reaction rates were reproduced (n=6, bovine liver catalase; n=3, 50v1 extract).

## Results

### Volume Displacement

The suitability of the VD apparatus for catalase enzymology was assessed by measuring the impacts of soil type, autoclaving, reaction time, substrate concentration, and repeated measurements on the rates of reaction. As exhibited in the rate plots in **Figure 1A**, addition of 330 mM hydrogen peroxide (1% *w/v* H_2_O_2_) to highly irrigated garden soils (CPP.BioTrek.Garden), biological soil crusts (Mojave.BSC.HD.LabStation), and permafrost (Alaskan.PF) resulted in the displacement of appreciable amounts of water at ~8, 4, and 0.2 g over 120 s, respectively. Autoclaved BSCs and permafrost provided no displacement (or activity), which supported a biochemical basis for the degradation of H_2_O_2_. Additionally, bare topsoils with very low ATP abundances, but appreciable 16S rRNA gene copy numbers (~10^5^ copy number g^−1^), provided no measurable rates by VD; thereby, indicating negligible contributions from geochemical sources (35).

**Figure 1.**
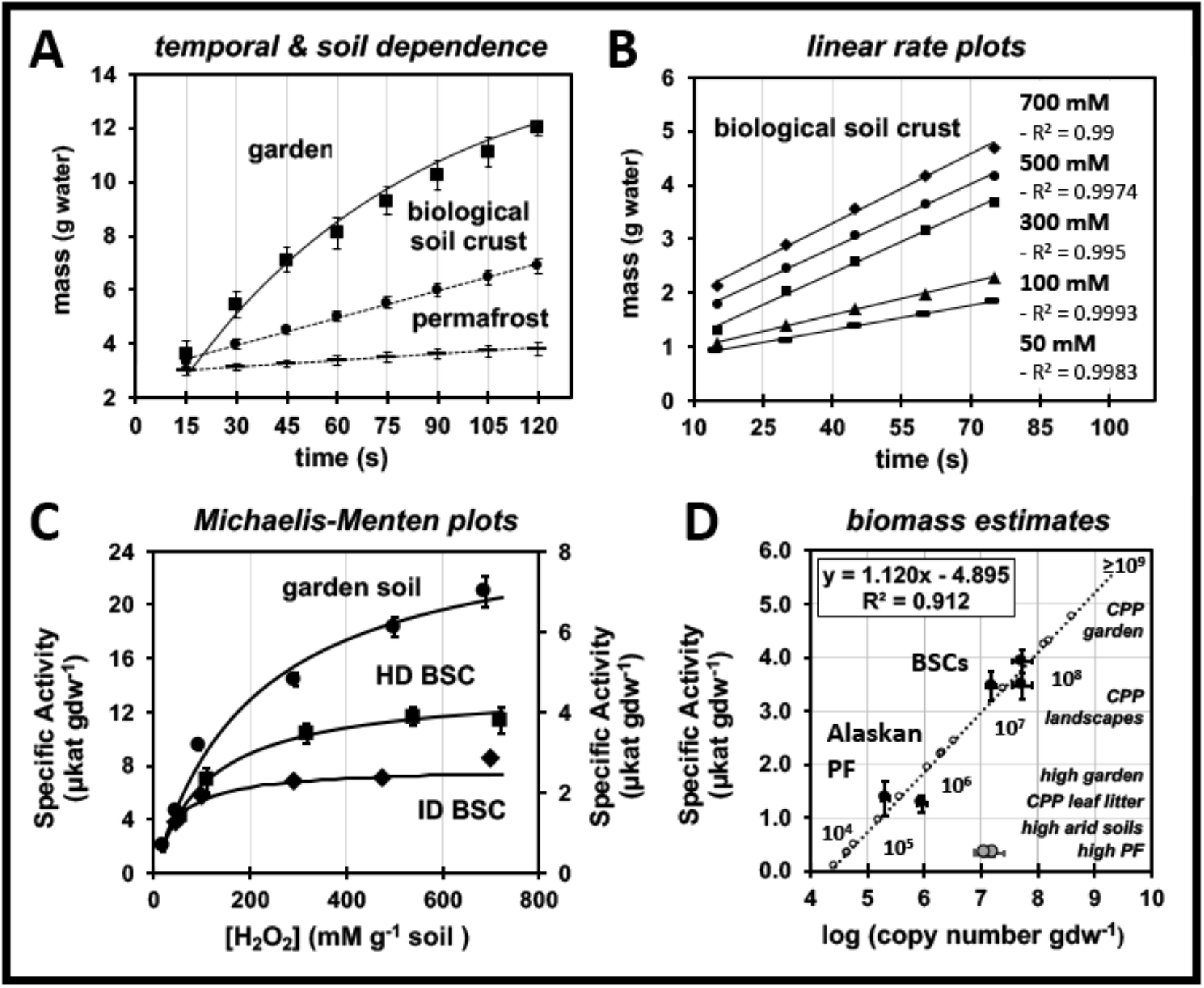
Kinetics and Michaelis-Menten analyses of soil catalase activities using volume displacement: (A) Temporal change in the mass of displaced water using highly irrigated garden soils (CPP.BioTrek.Garden), topsoils of biological soil crusts (BSCs obtained from areas of high surface density (Mojave.BSC.HD.Top), and 33 ky permafrost (Alaska.PF.33ky; 35k8C); errors bars represent the standard deviation (n = 3) and, for display purposes, trendlines for garden soils were estimated using psuedo-first order kinetics ([B]=[A]_0_(1-e^−*kt*^); where [A]_0_ and *k* were artificially set at 15 g and 0.014 s^−1^). (B) Impact of substrate concentration (50-700 mM) on the catalase activities for BSC topsoils, where linear regression of the rate plots from 15-75 s provided R^2^ ≥ 0.99. (C) Michaelis-Menten analyses for garden soils (left y-axis), and biological soil crusts (right y-axis) sampled from sites of high (HD) and intermediate (ID) surface densities; specific activities were expressed as μkat per gram dry weight (μkat gdw^−1^), errors bars represent the standard deviation (n=3), and fits to the data were obtained using **Equation 8**. (D) Standard curve (black squares; R^2^ = 0.91) representing relationship between catalase specific activities (μkat per gram dry weight; μkat gdw^−1^) and rRNA gene copy numbers (log copy number) using HD- and ID-BSCs (Mojave.BSC.HD.Top.CPP, Mojave.BSC.HD.Top.DSC, and Mojave.BSC.ID.Top.DSC), and 19 and 33 ky Alaskan permafrost (Alaska.PF.33ky and Alaska.PF.19ky); error bars along both axes represent the standard error (n=3), BSC subsurfaces are shown for comparison (gray circles), the 10^x^ labels represent probable biomass increments, and estimates of biomass (open circles) for Cal Poly Pomona (CPP) and Ladakh soils are listed (*italics*) in respective order.

Across the tested samples (3 tested soils; 3 technical replicates per soil type), a pooled relative standard deviation (RSD) of 5.1% for the measurements were obtained, which indicated a high repeatability for the VD technique. Reaction rates (g H_2_O displaced s^−1^) were calculated by linear regression with high certainty (R^2^ ≥ 0.99), with **Figure 1B** displaying representative examples for BSCs at differing substrate concentrations. As per **Figure 1C**, the change in catalase specific activities (μkat gdw^−1^) across 50-700 mM H_2_O_2_ for all tested soils reasonably conformed to regressions with the Michaelis-Menten equation (as supported by the residual sum of squares of 0.0412 for high-surface density (HD) BSCs, 0.0909 for intermediate-surface density (ID) BSCs, and 0.348 for the irrigated garden soil). In practice, the VD device (as assembled) exhibited a functional lower detection limit of ~0.2 μkat when using 1 g of soil (or ~12 Units), as was readily observed during analysis of alkaline evaporates, where catalase rates at 1-3 cm along the compressed core were not observable by VD, but were measurable by EC to provide apparent specific activities of ~0.09-0.14 μkat g^−1^. This limit was further reduced to 0.06 μkat g^−1^ when using 10 g of sample, as was observed with Ladakh permafrost.

### Impacts of Buffer & Substrate Formulation

The impacts of buffer identity and substrate formulation on the catalase specific activities (μkat gdw^−1^) across 7 different samples were measured by VD (**Figure S2**). Given the potential for inhibition by NaCl on catalases (36), we compared the impacts of 50 mM HEPES (pH 7.5) and 1x PBS on the enzyme reaction rates. In addition, we measured the inhibitory impacts of the stabilizing agents found in commercially-available H_2_O_2_ solutions. Formulations of H_2_O_2_ referred to as ‘stabilized H_2_O_2_’ contain chemical agents that prevent (or slow) the decomposition of H_2_O_2_ (*e.g.*, colloidal stannate, phosphates, sodium pyrophosphate, phosphonates, nitrate, and colloidal silicate (37, 38)). As per **Figure S2**, the highest catalase specific activities were obtained when using HEPES and non-stabilized H_2_O_2_. When using non-stabilized H_2_O_2_, comparisons across the buffers revealed an ~30-50% inhibition in PBS (p≤0.04) for microbial catalases from permafrost, moderately-irrigated landscaped soils with adjacent plant coverage (CPP.EnvDes.DryPatch), and dry soils with no adjacent plant coverage (CPP.BioTrek.DryPatch). When using HEPES, comparisons across H_2_O_2_ formulation revealed an ~30% inhibition when using stabilized H_2_O_2_ (p<0.02) for microbial catalases from BSCs (Mojave.BSC.Field) and moderately-irrigated landscaped soils soils with adjacent plant coverage (CPP.EnvDes.DryPatch).

### Comparison Across Techniques

Catalase SAs (**Figure 2**) for bovine liver catalase (BLC), a protein extract of *A. radioresistens* 50v1 (50v1), BSCs, and permafrost were measured by VD, EC, and/or absorbance spectroscopy (AS). Across these techniques, AS provided measures of substrate concentration (H_2_O_2_) in the aqueous phase, while VD and EC provided measures of product (O_2_) in the gas phase. To allow for comparisons across each of these techniques, SA values were expressed as the micromoles of H_2_O_2_ consumed per second per mg protein (μkat mg^−1^). For BLC, SA values from AS (339 ± 9 μkat mg^−1^) were ~5-fold higher than those from VD and EC (at 20 mM H_2_O_2_), which were equivalent (65 ± 12 μkat mg^−1^; 66 ± 19 μkat mg^−1^). Similarly, for the 50v1 extract, SA values from AS (6.3 ± 0.7 μkat mg^−1^) were ~3 and 6-fold higher than VD and EC, respectively (1.9 ± 0.1 μkat mg^−1^; 1.0 ± 0.3 μkat mg^−1^). This indicated that the degradation of H_2_O_2_ by soluble catalases resulted in speciation of O_2_ (over the timeframe used in regressions) as ~15-30% in the gas phase and ~70-85% dissolved in the aqueous buffer (and/or trapped as gas bubbles, which were typically visible after 30-80 s across all samples).

**Figure 2.**
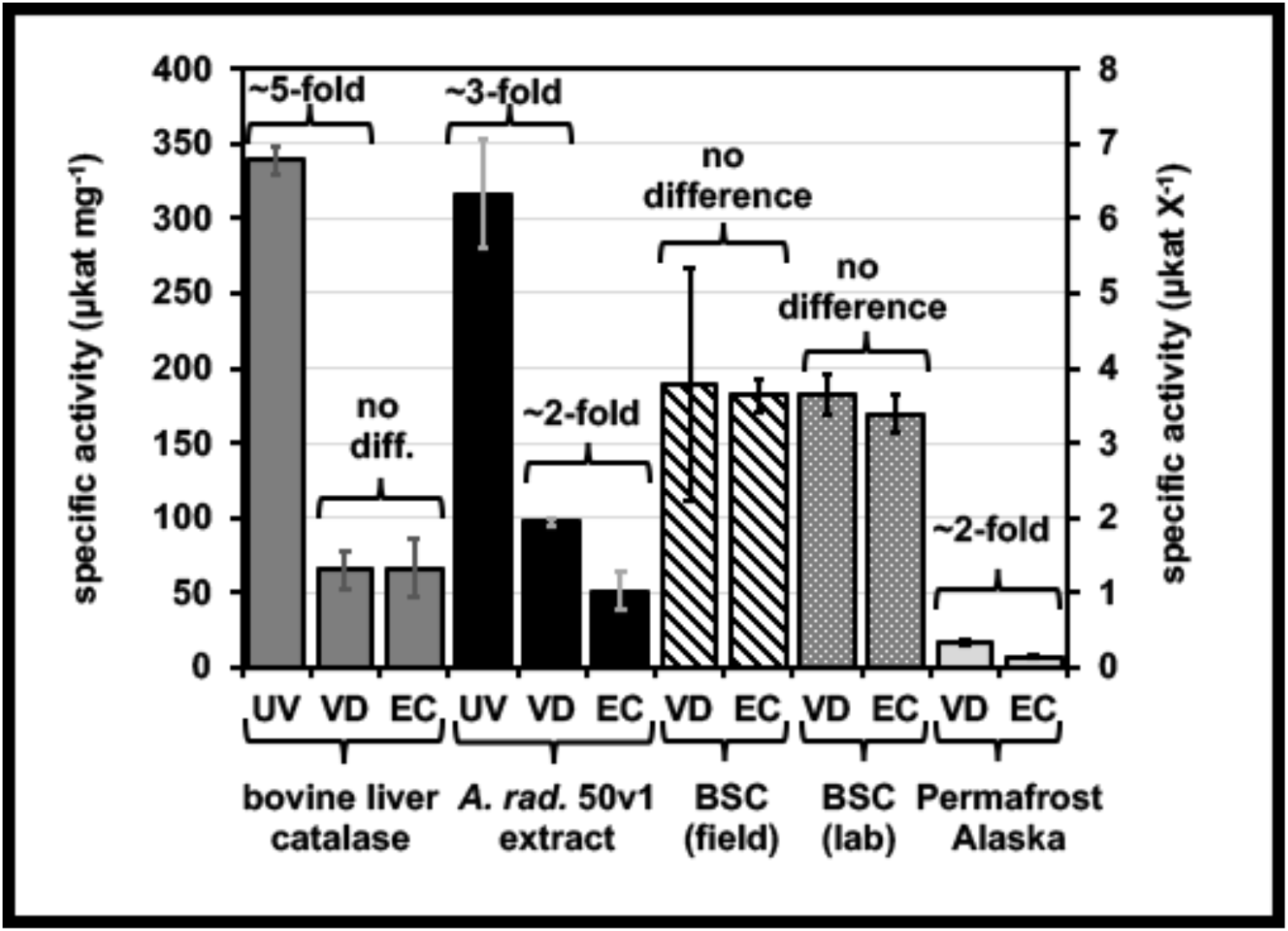
Comparison of catalase specific activities as measured by volume displacement (VD), electrochemical detection (EC), and ultraviolet absorption (UV), where specific activities are expressed as μkat mg^−1^ for bovine liver catalase (left y-axis) and as μkat X^−1^ (right y-axis) for a clarified extract of *A. radioresistens* 50v1 (X = mg), topsoils of biological soil crusts (BSCs) (X = g), and permafrost (X = g); error bars represent the standard deviation (n ≥ 3), and all assays were conducted in 50 mM HEPES (pH 7.5) using non-stabilized hydrogen peroxide, except for the BSC field assays which used stabilized hydrogen peroxide, and permafrost assays which used PBS.

For BSCs, traditional measures of catalase SAs were unsuccessful. When using AS, suspended soil particles provided substantial scatter in the absorbances. When titrating with KMnO_4_, BSCs provided uninterpretable data due to formation of foam in the reaction vessels (presumably due to the high organic content in the BSCs). Reproducible rates were only obtained when using VD or EC. Values for BSCs from VD and EC were respectively equivalent in field station and field experiments. In field station measurements, and when using non-stabilized H_2_O_2_, SAs of 4.0 ± 0.3 μkat gdw^−1^ (n=3) and 3.8 ± 0.3 μkat gdw^−1^ (n=5) were obtained from VD and EC, respectively. In *field*-based measurements, and when using stabilized H_2_O_2_, SAs of 4.3 ± 1.8 μkat gdw^−1^ (n=5) and 4.0 ± 0.2 μkat gdw^−1^ (n=2) were obtained from VD and EC, respectively. Together, this indicated that the moles of O_2_ estimated by volume displacement were equivalent to those measured by electrochemical detection (*e.g.*, O_2_ Gas Sensor). For the permafrost and 50v1 extract samples, however, VD reproducibly provided ~2-fold higher values than EC, which was suggestive of the presence of alternative gas-liberating degradation pathways for H_2_O_2_, which were observable by VD and AS, but not EC.

### Kinetic Model for Soil Catalases

Rate data were modeled to the catalase reaction sequence provided in **Diagram 2**, which begins with the soil matrix catalases (*E*_*soil*_) in the reduced state, inclusive of the Fe^II^(heme) and (Mn^II^)_2_ cofactors; catalysis beginning with the oxidized Mn catalases was considered to be minimal (20). Therefore, in this model the combined steady state included formation and breakdown of the oxidized soil microbial catalases 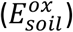, inclusive of Compound 1 and the (Mn^III^)_2_ catalase (**Equations 5 & 6**) (13, 14, 39).

As described by *k*_1_ (M^−1^ s^−1^), the formation of 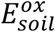 encompassed several steps including equilibration of H_2_O_2_ into the soil suspension, capture of H_2_O_2_ by the catalases (*E*_*soil*_) in the biological matrix, and irreversible formation of the oxidized enzyme. As per *k*_R_ (s^−1^), non-productive breakdown of 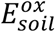 included degradation back to native enzyme via Compound II and/or reduction of the Mn cofactor. Productive breakdown of 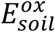, as per 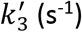, was treated as a pseudo 1^st^ order reaction (where 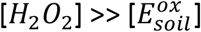) that encompassed rapid acquisition of the second substrate (H_2_O_2_), formation of products (O_2_ and H_2_O), and release of O_2_ into the aqueous phase. Lastly, equilibrium of O_2_ into the gas phase (thereby allowing detection by VD and EC) was described by Henry’s Law 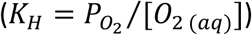.

**Diagram 2.**
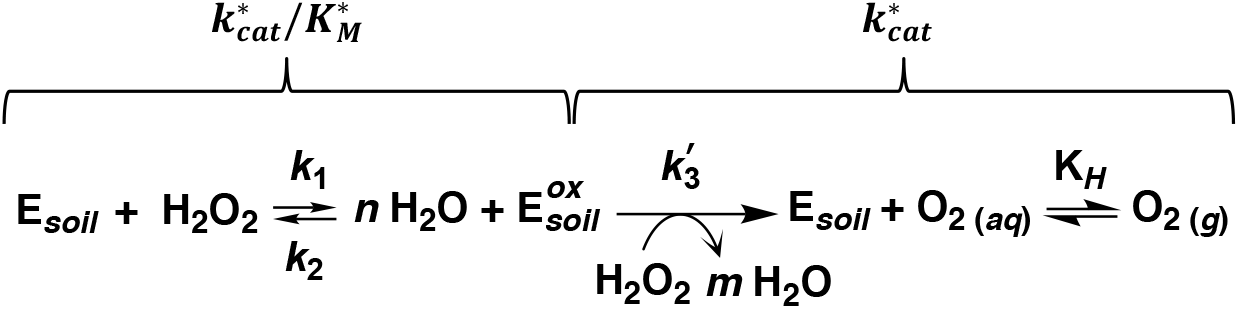
Simplified reaction sequence for soil catalases (*E*_*soil*_), where n=1 and m=1 for Fe(heme)-catalases, and n=2 and m=0 for Mn-catalases.

As expressed in **Equation 7**, the rate equation for product formation (or detection) included enzymatic formation 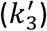 and liberation of O_2_ into the gas phase. As per Henry’s Law, the moles of gaseous O_2_ were obtained from the oxygen partial pressure and ideal gas law (*K*_*H*_ = *n O*_2(*g*)_ (*RT*/*V*)/[*O*_2 (*aq*)_]). In turn, a modified Michaelis-Menten equation (**Equation 8**) was derived by inclusion of steady state and mass balance terms (**Equation 6**), where 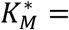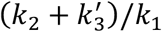. To incorporate soil mass terms, a soil biomass ratio (*R*_*s*_) was introduced into the kinetic treatment, where *R*_*s*_ related the total moles of catalase in the soil microbial community to the equivalent and total grams of dried bioactive soil 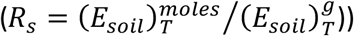. Thus, as per **Equation 6**, the total enzyme concentration, [*E*_*soil*_]_*T*_, was expressed as 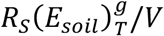. For this expression, multiplication of *R*_*s*_ by 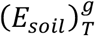 yielded 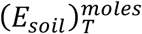, which in turn provided [*E*_*soil*_]_*T*_ after division by the reaction volume (*V*). As per **Equation 8**, the assembled rate equation was thus simplified by combining 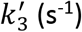, *K*_*H*_*V*/*RT* (L), and *R*_*s*_/*V* (M gdw^−1^) to yield the parameter of 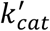 (in units of moles s^−1^ gdw^−1^).

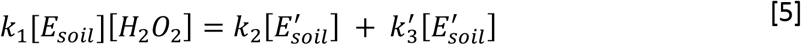

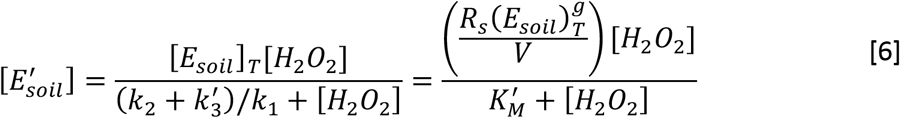

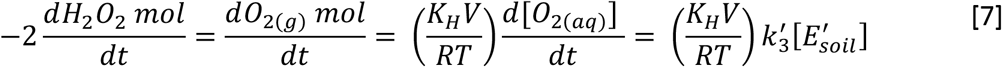

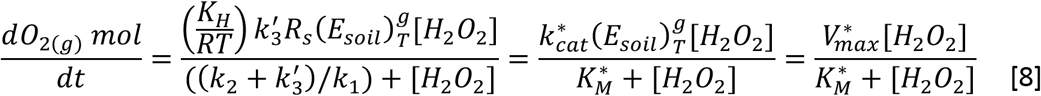

Accordingly, regression analyses provided the *apparent* (*) terms of 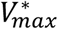 (μmole s^−1^; μkat) and 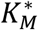 (mM g^−1^), which represented the maximal rate of H_2_O_2_ degradation by the soil catalase community 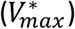, and the [H_2_O_2_] required (per gram of soil) to reach 50% of the 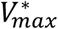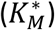. For this study, the calculated term of *apparent* turnover number 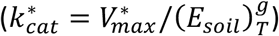 represented the maximal rate of H_2_O_2_ degradation per gram of dried soil (μkat gdw^−1^), and the calculated term of *apparent* specificity constant 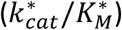 was expressed in units of gdw^−1^ s^−1^ (by converting 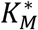 to moles of H_2_O_2_ using the reaction volume and 1 g of soil).

Under the described experimental conditions, the rate limiting steps among the productive reactions were presumed to be substrate capture by the soil catalases (a component of *k*_1_) and/or release from the soil matrix (a component of 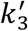). Non-productive degradation of 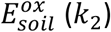 (*k*_2_) was assumed to be a minor reaction component, where 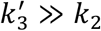 (40–42). Therefore, under these total assumptions, the 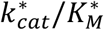 term effectively reduced to (*K*_*H*_/*RT*)*k*_1_*R*_*s*_, and was expressed in simplified units of gdw^−1^ s^−1^ (through division by the reaction volume). In effect, these units were consistent with a rate constant for a second order reaction – albeit, in mass-based terms. Accordingly, the 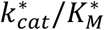 term (~*k*_1_) was interpreted as relating to the rates of substrate capture by the catalases (**Diagram 2**), which included acquisition of H_2_O_2_ and the first irreversible step of catalysis (formation of 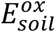). Similarly, the 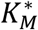 term reduced to _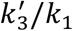_ (mM g^−1^), or a ratio of the rates of product release over substrate capture, and was interpreted as the capacity of the soil to degrade H_2_O_2_. Thus, as per **Diagram 2**, the 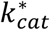 term was interpreted as relating to the rates of gaseous O_2_ release.

### Michaelis-Menten Kinetics

Michaelis-Menten kinetic studies were performed on (1) topsoils of BSCs obtained from areas of high (HD) and intermediate (ID) surface coverages (HD.Mojave.BSC.LabStation & ID.Mojave.BSC.LabStation, respectively), and (2) highly-irrigated CPP gardens soils (CPP.BioTrek.Garden). Across the 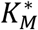 values (**Figure 3A**), CPP garden soils (210 ± 40 mM) exhibited an ~2 and 5-fold greater capacity to degrade H_2_O_2_, per gram of soil, as compared to the BSCs (110 ± 15 and 45 ± 13 mM). Across the BSC samples, the HD-BSCs exhibited an ~2-fold greater capacity to degrade H_2_O_2_ per gram of soil (p<0.05) than the ID-BSCs.

**Figure 3.**
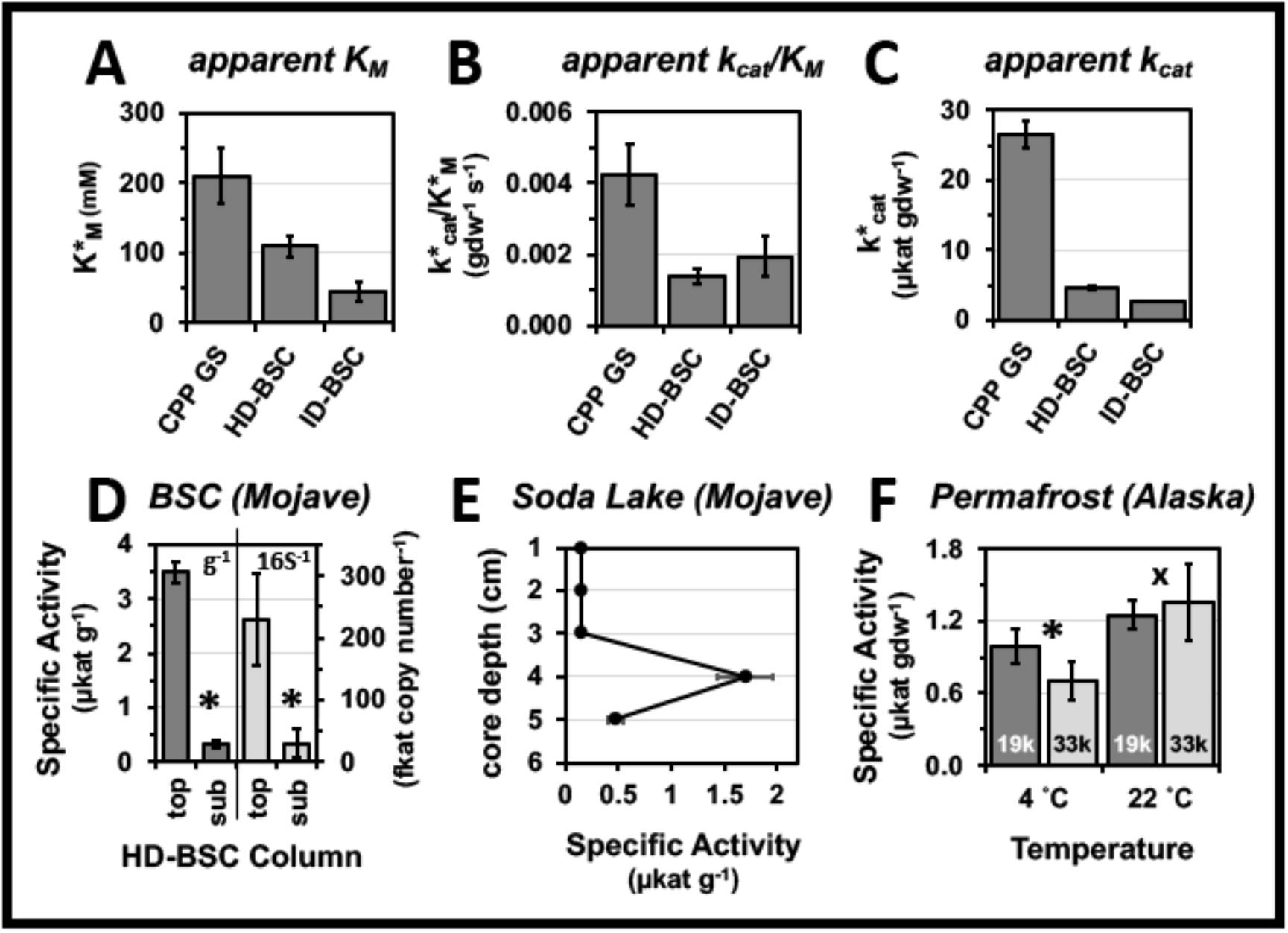
Trends across soil catalase kinetics and specific activities where statistical comparisons are marked (*p < 0.05; ^X^p > 0.05): Comparison of the Michaelis-Menten kinetic parameters for (A) 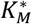, (B) 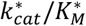, (C) and 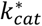 across highly irrigated garden soils (CPP GS; CPP.BioTrek.Garden), and topsoils of biological soil crusts (BSCs) sampled from sites of high (HD-BSC; BSC.HD.Mojave.Lab) and intermediate (ID-BSC; BSC.ID.Mojave.Lab) surface densities; rates for CPP GS and HD-BSC were measured in triplicate (n=1 for ID-BSC), and error bars represent the standard error of the regression. (D) Comparison of specific activities for HD-BSCs along the vertical column structure (topsoil vs. subsurface = top vs. sub) where specific activities are expressed as μkat per total soil mass (left y-axis; μkat g^−1^) and fkat per 16S rRNA gene copy number (right y-axis; fkat copy number^−1^); error bars represent the propagated error (n=3). (E) Change in specific activities (μkat g^−1^) along a depth profile for a compressed dry lake bed core; error bars represent the standard deviation (n=2). (F) Comparison of specific activities of 19 ky and 33 ky permafrost (Alaska.PF.33ky & Alaska.PF.19ky) measured at 4 and 22 °C, and expressed as μkat gdw^−1^; error bars represent the standard deviation (n≥4).

As displayed in **Figure 3B**, 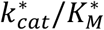 values for CPP garden soils (4.2×10^−3^ ± 0.9×10^−3^ gdw^−1^ s^−1^) were ~2-3-fold higher than those of BSCs (1.4×10^−3^ ± 0.2×10^−3^ and 1.9×10^−3^ ± 0.6×10^−3^ gdw^−1^ s^−1^); while values across the BSCs were statistically equivalent (p>0.05). This indicated that CPP garden soils exhibited the largest rates of substrate capture, whereas the rates across the BSCs were equivalent. Trends across the 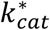 values (**Figure 3C**) indicated that CPP garden soils (27 ± 2 μkat gdw^−1^) were ~6-10-fold higher than the BSCs (4.6 ± 0.2 and 2.6 ± 0.2 μkat gdw^−1^). Across the BSCs, 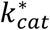 values for HD BSCs were ~2-fold higher (p<0.05) than those for ID BSCs. This indicated that CPP gardens soils exhibited the largest rates of product release. Across the BSCs, HD-BSCs displayed the largest rates of product release.

### Catalase Specific Activities

Catalase SAs across several types of soil microbial communities were measured by VD (**Figures 4** and **S2**, **Table S1**); including BSCs, Alaskan permafrost, high-elevation permafrost, high-elevation arid soils, high-elevation garden soils, temperate garden soils, and temperate landscaped soils. Reaction conditions included 50 mM HEPES (pH 7.5) with 330 mM non-stabilized H_2_O_2_, or 1x PBS with 330 mM stabilized H_2_O_2_; respectively referred to herein as HPS/NS and PBS/S. Trends across the SAs in HPS/NS (**Figures 4** and **S2**) were as follows: CPP garden soils (10 ± 1 μkat gdw^−1^) > BSCs measured in the field (8.4 ± 0.4 μkat gdw^−1^) > dry landscaped CPP soils (~5-6 μkat gdw^−1^) > BSCs measured in field station and formal laboratories (~3-4 μkat gdw^−1^) ≈ arid CPP soils (~ 3.4 ± 0.2 μkat gdw^−1^) > CPP garden soils under leaf cover (1.9 ± 0.1 μkat gdw^−1^) > Alaskan permafrost (1.2 ± 0.1 μkat gdw^−1^).

**Figure 4.**
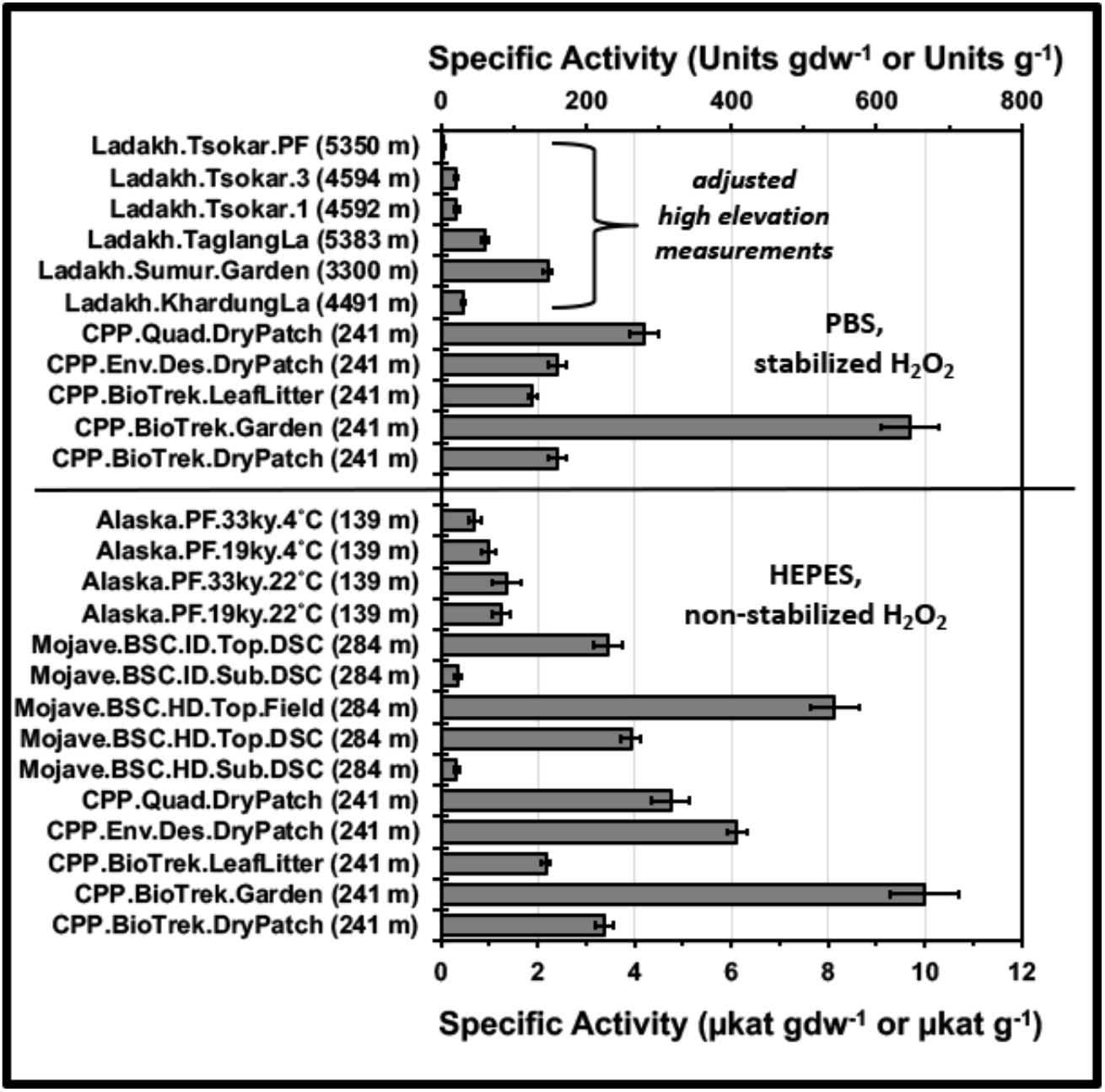
Comparison of barometrically adjusted catalase specific activities across differing soils (**Table S1**) expressed in SI units of μkat per gram dry weight (bottom x-axis; μkat gdw^−1^) and typical enzymology terms of Units per gram dry weight (top x-axis; Units gdw^−1^), with all Ladakh samples being expressed per total soil mass (g^−1^); measurements listed in the top panel were obtained in 1x PBS and 330 mM (1% w/w) stabilized hydrogen peroxide, measurements listed in the bottom panel were obtained in 50 mM HEPES (pH 7.5) and 330 mM (1% w/w) non-stabilized hydrogen peroxide, all error bars represent the standard deviation (n ≥ 3), sampling site elevations are given in parentheses, and elevations for measurements were 241 m (CPP.X soils, Mojave.HD.BSC.Top.CPP, & Alaska.PF.X), 284 m (Mojave.BSC.HD.Top.Field, Mojave.BSC.HD/ID.Top.DSC, & Mojave.BSC.HD.Top/Sub.DSC), and 3300 m (Ladakh.X soils).

For the BSCs, the highest SAs were obtained from field samples analyzed immediately after collection (7.2 ± 0.4 μkat g^−1^ or 8.4 ± 0.4 μkat gdw^−1^). After storage for ~2 days, SAs were ~2-fold lower (3.5 ± 0.2 μkat g^−1^), which indicated post-sampling degradation. After ~2 weeks of storage, measurements (in a formal laboratory) showed minimal further degradation (3.1 ± 0.2 μkat g^−1^). The SAs for BSCs were additionally compared along the vertical column structure (**Figure 3D**), with the topsoils (3.5 ± 0.2 μkat g^−1^) having ~10-fold higher SA values than the subsurface (0.33 ± 0.05 μkat g^−1^).

Microbial communities from high-elevation topsoils (Ladakh, India; 3300-5400 m) exhibited relatively lower SAs of ≤2.5 μkat g^−1^ (**Figure 4**). The lowest values were obtained from arid topsoils, which exhibited a range of ~0.4-0.8 μkat g^−1^ (Tsokar, Khardung La, and Taglang La), and from permafrost samples (Tsokar), which exhibited 0.6 ± 0.3 μkat g^−1^. Garden soils (Ladakh.Sumur) exhibited the highest value of 2.2 ± 0.1 μkat g^−1^. Samples obtained from the slightly alkaline Panamik hot springs (pH ~8) yielded no VD rates (when at <25 °C), likely due to inhibition by sulfide (43), despite the clear visual evidence of microbial mats. For the alkaline evaporates from Soda lake in the Mojave National Preserve (**Figure 3E**), SAs increased ~12-fold between 1-3 cm (~0.09-0.14 μkat g^−1^) and 4 cm (1.7 ± 0.3 μkat g^−1^) along the compressed core, and then decreased ~4-fold at 5 cm (0.47 ± 0.07 μkat g^−1^).

For Alaskan permafrost (**Figure 3F**), SAs were measured at 4 and 22 °C for samples collected across a chronosequence of 19 and 33 ky before present. At 4 °C, SAs (HPS/NS) for the 19 ky sample (0.99 ± 0.14 μkat gdw^−1^) were ~1.4-fold higher than the 33 ky sample (0.70 ± 0.12 μkat gdw^−1^). At 22 °C, the SAs were statistically equivalent (~1.2-1.4 μkat gdw^−1^). For the 19 ky samples, SAs were similar across the temperatures (0.99 ± 0.15 & 1.2 ± 0.16 μkat gdw^−1^); however, for the 33 ky sample, SAs were higher at 22 °C by a factor of 1.9 ± 0.6 (p = 0.019).

### Standard Curve for Biomass Estimation

Estimates of microbial biomass in soils and permafrost were obtained from a standard curve (**Figure 1D**), which was assembled using catalase SAs (μkat gdw^−1^) and measured 16S rRNA gene copy numbers for HD-BSC topsoils (5.3×10^7^ ± 2.2×10^7^ copy number gdw^−1^), ID-BSC topsoils (1.5×10^7^ ± 0.3×10^7^ copy number gdw^−1^), 19 ky Alaskan permafrost (9.0×10^5^ ± 1.7×10^5^ copy number gdw^−1^), and 33 ky Alaskan permafrost (2.0×10^5^ ± 0.1×10^5^ copy number gdw^−1^) (35, 44). For this study, bacterial rRNA gene copy numbers were used as a proxy for biomass (as eukaryotic (18S) rRNA gene copy numbers for these samples had yet to be determined). Linear regression across the standard curve provided good fits (R^2^ = 0.91), respective standard errors of 18 and 28% for the slope and intercept, and 1.1×10^4^ 16S rRNA gene copy number gdw^−1^ as the limit of detection (LOD).

Estimates of biomass (copy number gdw^−1^) for the soil microbial communities across the CPP campus were ~10^13^ for garden soils (BioTrek.Garden), ~10^8-9^ for moderately-irrigated landscaped soils (CPP.Quad.DryPatch & CPP.EnvDes.DryPatch), ~10^7^ for irregularly-irrigated soils (CPP.BioTrek.DryPatch), and ~10^6^ for garden soils under substantial leaf cover. Across the microbial communities from high-elevation soils, biomass estimates (copy number gdw^−1^) were ~10^6-9^ for garden soils (when assuming soil water contents of ~70% or less; Ladakh.Sumur), ~10^4-5^ for arid topsoils (Ladakh.Tsokar, Ladakh.KhardungLa, and Ladakh.TaglangLa), and ~10^4^ for permafrost (Ladakh.Tsokar.PF). Bacterial abundances for garden soils were potentially high due to exclusion of eukaryotic biomass.

Due to experimental variances across the samples (*e.g.*, use of HPS/NS, PBS/S, dry soil mass, and total soil mass), biomasses were only estimated in ~10^x^ increments (which amounted to differences of ≥1.1 μkat gdw^−1^ between the estimates). In addition, biomasses were only estimated using SAs obtained from field station or formal laboratory measurements (similar to the standard curve). Catalase SAs for BSC subsurfaces (**Figure 1D**) did not follow the regression trend (measured biomasses for HD-BSCs and ID-BSCs were 1.5×10^7^ ± 0.8×10^7^ and 1.2×10^7^ ± 0.5×10^7^ copy number g^−1^, respectively (35)). As a result, the standard curve and biomass estimates were restricted to topsoils and permafrost; or, to environmental samples that were likely accustomed to appreciable native oxidative stresses prior to collection (as was assumed/inferred for the topsoils due to continual aerobic and photosynthetic metabolism and exposures to ultraviolet radiation; and for permafrost due to metagenomic and genomic lines of evidence which show high abundances of genes associated with oxidative stress (45–47)).

## Discussion

### Portable Devices and Field-Amenable Assays

Low-cost, portable, and field-amenable devices were assembled to measure microbial catalase activities in soils and permafrost. Portability and field-applicability for VD were readily demonstrated through experiments conducted in the field (Mojave National Preserve), temporary work station (Mogol Hostel, Ladakh, India), and field station laboratory (CSU Desert Studies Center). Given the need for continual upright storage of the O_2_ Gas Sensor for EC, the VD apparatus was better suited for field campaigns and travel. Measurements by VD were rapid (<2 min per sample) and amenable to the analysis of large samples sets. In contrast, measurements by EC exhibited appreciable lag times in the reactions, and required re-equilibration of the sensor between runs. Nevertheless, for lower activity samples (<0.2 μkat, <12 Units; 1-2 g sample), EC was more reliable, as was demonstrated with alkaline evaporate samples (however, a geochemical basis for the low degradation rates could not be ruled out).

To enact comparisons across measurements obtained during field campaigns, rates were barometrically adjusted to correct for the impacts of elevation, relative humidity, and temperature. For catalase SAs from high-elevation soil microbial communities, this prevented over-inflation by ~1.5-fold. To afford comparisons across spectral, electrochemical, and physical techniques, displacement rates (g H_2_O displaced s^−1^) and electrochemical rates (%O_2_ min^−1^) were transformed to molecular rates using SI units (μkat, or μmoles H_2_O_2_ consumed s^−1^). Such conversions are atypical for environmental catalases.

Equivalent catalase SAs (p>0.05) were obtained from VD and EC when using BSCs (in lab station and field experiments); thereby directly correlating the displacement of water to the electrochemical detection of gaseous O_2_. For permafrost and bacterial samples, the ~2-fold higher values from VD were suggestive of the presence of gaseous side products (*e.g.*, CO_2_ and H_2_) potentially formed from reactions (*e.g.*, oxidation and homolytic fragmentations) between H_2_O_2_ and cellular carbohydrates (48). While speculative, these results were suggestive of extra/intracellular carbohydrates from ice-laden permafrost and aqueous bacterial cultures (50v1 extract) being more prone to degradation by H_2_O_2_ than those from the desert microbial communities (BSCs).

Across the tested buffers and substrates, the trends suggested that microbial catalases arising from nutrient-limited or stressed environments (*e.g.*, permafrost, BSCs, and dry soils) were susceptible to inhibition by the H_2_O_2_ stabilizing agents and/or higher-ionic strength solutions. This indicated that optimal reaction conditions for comparison of microbial catalases from extreme environments were 50 mM HEPES (pH 7.5) and ≥300 mM non-stabilized H_2_O_2_. As a drawback, however, these conditions necessitated cold storage (at ≤4°C) for non-stabilized H_2_O_2_ to maintain reagent integrity. Given this impracticality for field campaigns, alternative conditions were 1x PBS and ≥300 mM stabilized H_2_O_2_; with advantages including ease of buffer preparation (*e.g.*, dissolution of tablets, or dilution of commercially available 10x PBS solutions), and the long-term ambient stability and commercial availability of 3% stabilized H_2_O_2_.

### Kinetics of Soil Microbial Catalases

Catalase kinetics were measured in suspensions of BSCs and gardens soils. Across the samples, CPP garden soils exhibited the highest capacity to degrade H_2_O_2_ 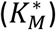, fastest rate of substrate capture 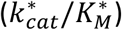, and fastest rate of gaseous product release 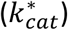. As per our soil catalase model, these trends were likely the result of high microbial abundances, rather than major differences in the intrinsic rate constants (for the catalase community). As per **Equation 8**, high biomass values (*e.g.*, high cells per gdw) propagate through *R*_*s*_ (as moles of catalase per gdw) and directly raise the 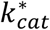 and 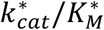 terms, since 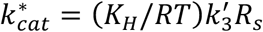. To obtain upper estimates of *R*_*s*_, therefore, we assumed that (1) 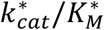 was approximately equal to (*K*_*H*_/*RT*)*k*_1_*R*_*s*_ when 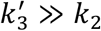, (2) values for *K*_*H*_ (756 atm M^−1^) were roughly similar across the soil suspensions, (3) rate constants for bimolecular reactions in the soil/biological suspensions were reduced ~100-fold due to viscosity changes (49), and (4) commensurate ~100-fold reductions in the representative and diffusion-limited *k*_*cat*_/*K*_*M*_ term (~10^7^ M^−1^ s^−1^) for soluble catalases (in classical Michaelis-Menten kinetics, *k_cat_*/*K*_*M*_ ≈ *k*_1_ when *k*_2_ ≫ *k*_−1_).

With these assumptions, we respectively obtained *R*_*s*_ values of ~40, 13, and 19 pmol gdw^−1^ for CPP garden soils, HD-BSCs, and ID-BSCs, which indicated >2-fold higher catalase abundances per soil mass for the garden soils. In turn, accounting for *R*_*s*_ within the 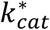 term provided estimates of 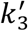. Respective estimates for 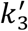 were ~2.1×10^4^, 1.1×10^4^, and 0.45×10^4^ s^−1^ for CPP garden soils, HD-BSCs, and ID-BSCs. These rate constants were consistent with the reported *K*_i)5_ values (~10^3-6^ s^−1^) for purified catalases (10, 36, 50, 51); thereby providing support to the above assumptions. The trends across the rate constants also implied that the ~2-fold larger 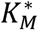 values 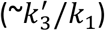 for CPP garden soils were not due to major changes in substrate acquisition, but rather the result of a ~2-fold higher rate of product formation 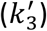.

When comparing the HD and ID-BSCs, the HD-BSCs exhibited a higher capacity to degrade H_2_O_2_ 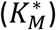, an equivalent rate of substrate capture 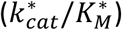, and faster rate of gaseous product release 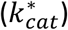. The similar 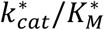 values supported the rates of substrate capture (*k*_1_) being diffusion-limited in the soil suspensions (as was assumed in the calculation of *R*_*s*_). Trends across the rate constants again implied that the ~2-fold larger 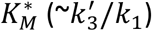 for HD-BSCs was the result of the ~2-fold higher rates of product formation 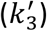.

### Trends in Catalase Specific Activities

Comparisons across differing soils and permafrost supported the use of catalase SAs (**Figure 4**) as markers for microbial biomass and indicators for biological activity. For microbial communities from arid soils, catalase SAs per soil mass (μkat gdw^−1^, μkat g^−1^) from high elevation Ladakh topsoils (Tsokar, Khardung La, and Taglang La; 4500-5500 m) were ~10-fold lower than those from BSC topsoils, which were sampled at much lower elevations (450-685 m). As per our model, these differences corresponded to lower microbial biomasses of ~10^4-5^ copy number gdw^−1^ for Ladakh topsoils (calculated value), as compared to ~10^7^ copy number gdw^−1^ for BSCs (measured value). For permafrost, catalase SAs obtained at 4 and 22 °C amounted to activation energies of 9-25 kJ/mol, which was consistent with the literature values for purified catalases (39, 51). Catalase SAs (0.06-1.4 μkat gdw^−1^ or μkat g^−1^) were also measurable in samples arising from anaerobic and/or oxygen-limited environments, including BSC subsurfaces, Alaskan permafrost, Ladakh permafrost, and Soda lake alkaline evaporates. Hence, these results suggested that the anaerobic and/or oxygen-tolerant microbial communities experience oxidative stress due to the presence of active catalase enzymes.

### Catalases and Microbial Biomass

To obtain further insights into the relationships between catalase activity and biomass, the values of *R*_*s*_ and SA were normalized to 16S rRNA gene copy numbers per gram dry weight (copy number gdw^−1^). Conversions were conducted using measured 16S rRNA gene copy numbers for BSC topsoils, BSC subsurfaces, and Alaskan permafrost; while calculated values were used for CPP and Ladakh topsoils. For *R*_*s*_, re-expression provided catalase abundances of ~250 and 1200 zeptomoles (zmol) per 16S rRNA gene copy number for HD- and ID-BSCs, and 0.004 to 4 zmol copy number^−1^ for CPP garden soils (when assuming 1×10^10-13^ copy number gdw^−1^). Considering an average of two 16S rRNA gene copy numbers per cell (52, 53), and a representative cellular volume of ~1 μm^3^ for soil microbes (54, 55), these values amounted to catalase concentrations of ~0.5 mM for HD-BSCs, ~2 mM for ID-BSCs, and ~0.008-8 μM for CPP garden soils. In context, cultured *E. coli* cells (~0.6 μm^3^) reportedly contain ~0.03 mM catalase when exposed to mild external oxidative stress (100 μM exogenous H_2_O_2_), and up to ~0.4 mM when subjective to moderate intracellular oxidative stress (50 μM intracellular H_2_O_2_) (56–58). Therefore, consistent with these comparisons is the assessment that microbial communities from BSCs experience substantially higher degrees of native oxidative stress and, consequently, possess substantially higher basal concentrations of intracellular catalase (≤10^5^-fold higher concentrations than garden soils communities, as per these estimates).

In support of this assessment are catalase SAs expressed per biomass, which also increased for communities potentially subjected to high degrees of native oxidative stress. As per **Figure 5**, trends for SAs per biomass (fkat copy number^−1^) across the samples were 33 ky Alaskan permafrost (6700) > high-elevation Ladakh topsoils (5900–6600) > Ladakh permafrost (1900) > 19 ky Alaskan permafrost (1400) = CPP leaf litter soil (1400) > Sumur garden soils (1300) > dry CPP soils (250) > HD- and ID-BSC topsoils (74, 230) > HD- and ID-BSC subsurfaces (22, 30) > CPP landscaped soils (~3-30) > CPP garden soils (0.004). These trends suggested that microbial communities in permafrost, high-elevation topsoils, and topsoils under leaf litter experienced appreciable oxidative stress. For permafrost communities, native stresses potentially included background radiation dosages accumulated over geological time scales (59, 60). For high-elevation arid topsoil communities, stresses included the accumulated dosages from intense UV exposures, where UV-A doses are ~10x greater than sea level (33). And, for topsoil communities under leaf litter, stresses included persistent exposures to H_2_O_2_ (and other reactive oxygen species) formed during plant-matter degradation (61–63).

**Figure 5.**
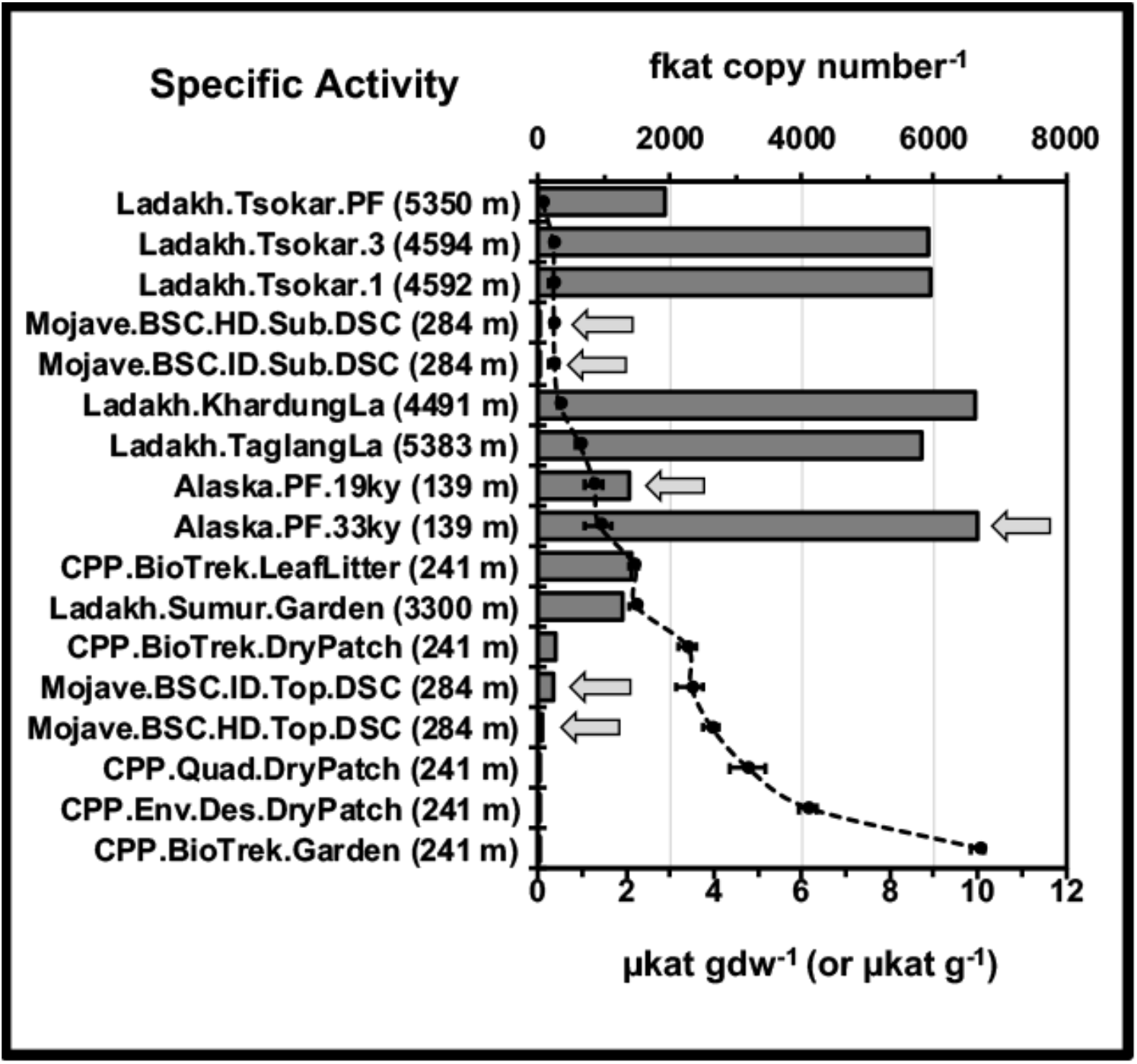
Comparison of specific activity (SA) values expressed per biomass (top x-axis; bars) and soil mass (bottom x-axis; black circles), where SAs per biomass are expressed in units of fkat per 16S rRNA gene copy number (fkat copy number^−1^) and SAs per soil mass are expressed in units of μkat per gram dry weight (μkat gdw^−1^), or per total total mass for the Ladakh samples (μkat g^−1^); soil samples (y-axis) are listed in order of increasing SA per soil mass (top to bottom), arrows denote measured 16S rRNA copy number values (with all others being calculated), dotted line represents the trend, and error bars represent the standard deviation for the specific activities per soil mass along the x-axis (μkat gdw^−1^, or g^−1^; n≥3).

In comparison, the lowest SAs obtained from garden and landscaped soils were suggestive of minimal degrees of native oxidative stress for high biomass microbial communities from regularly irrigated soils. Further, for black-crusted BSCs, catalase SAs revealed differing degrees of stress along the vertical column structure (**Figure 3D**); where representative catalase SAs per biomass were ~8-fold higher in the topsoils (230 ± 74 fkat copy number^−1^) than the subsurfaces (30 ± 23 fkat copy number^−1^). Potential sources of reactive oxygen species for BSC topsoil communities included ultraviolet radiation exposures, photosynthesis, and aerobic respiration (4–6).

### Implications for Microbial Ecology

In the context of microbial ecology, the differing profiles across 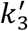, *R*_*s*_ per biomass, and SAs per biomass served as indicators for changes in catalase community structure (*e.g*, differences in 1° structure, metal cofactor, and class) and, by extension, microbial community structure. For instance, when comparing HD-BSCs to CPP garden soils, the ~2-fold lower values for 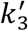, ≥10^5^-fold higher values for *R*_*s*_ per biomass, and ~10^4^-fold higher SAs per biomass were consistent with broad phylum-level taxonomic changes in the community structure. In support, phylogenetic studies show that HD-BSCs are numerically dominated by Cyanobacteria and Proteobacteria (35), while garden soils are typically dominated by Actinobacteria, Proteobacteria, and Firmicutes (64, 65). When comparing HD-BSCs to ID-BSCs, however, the mild differences in kinetics (*e.g.* ~2-fold higher values for 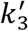, ~5-fold lower values for *R*_*s*_ per biomass, and ~3-fold lower SAs per biomass) were consistent with moderate genus-level changes across the Cyanobacteria and Proteobacteria, as phylum distributions were equivalent in these samples (35). In support, comparisons conducted for this report sBhSoCws ctohnattatihne~1H.D5-fold lower abundances of *Phormidium*, ~1.7-fold higher abundances of an unidentified genus from *Nostocophycideae*, and a ~2-fold lower abundance of an unidentified genus from *Oxalobacteraceae* (when using a Benjamini–Hochberg critical value of *p* < 0.0099, false discovery rate of 0.50). Similarly, for Alaskan permafrost, the ~5-fold increase in catalase SAs per biomass across the chronosequence (19 vs. 33 ky samples) were attributed to the presence of cold-adapted catalases arising from broad phylum-level changes in community structure with Acidobacteria, Proteobacteria, and Bacteroidetes dominating the 19 ky samples, and Firmicutes dominating the 33 ky samples (46).

## Conclusions

In conclusion, the combined kinetic trends described in this report support the hypothesis that microbial communities which experience higher degrees of oxidative stress possess higher basal concentrations of intracellular catalase (*R*_*s*_ per copy number) and catalase specific activities per biomass (SAs per copy number), and that differing kinetic profiles across catalase communities are indicative of phylogenetic changes in community structure. In addition, we aptly demonstrate that volume displacement serves as a cost-effective, simple, and field-amenable method for measuring catalase kinetics across differing environmental samples. Further, this method is suitable for scientists and educators from all disciplines, irrespective of budgetary concerns, or familiarity with chemical kinetics. Thus, as a biochemical tool for microbial ecology, this assay and kinetic treatment represents a robust means to detect and quantify the presence and abundance of active microbial communities in soils and permafrost.

## Author Contributions

All authors contributed to the acquisition and analysis of the data. Contributions to critical reports/revisions of the work were obtained from MC, EM, JC, NR, SG, and RM; and all contributors approved the manuscript, and agreed to be accountable for the work. The primary investigator and corresponding author is RM. Michaelis-Menten analyses, specific activity measurements of BSCs, comparative controls, and development of barometric calculations were performed by MC. Critical initial measurements and field work with VD were conducted by EM. Comparisons of buffers and substrate formulations on the specific activities of CPP soils were conducted by JC and NR. Permafrost measurements by VD and EC were conducted by SG.

## Acknowledgments

This work was funded through the NASA Astrobiology Minority Institutional Research Support program (MIRS), and the California State University (CSU) Math and Science Teacher Initiative (MSTI). We acknowledge the contributions of Nicholas Cooper who performed field-based measurements of BSCs by EC, as well comparative laboratory controls. Catalase-based field work was conducted as part of the NASA/CSU Spaceward Bound pre-service teacher training program in the Mojave National Preserve (2016), and as part of a Spaceward Bound field campaign in Ladakh, India (2017). Scientific Research and Collecting Permits from the National Park Service for the Mojave National Preserve included MOJA-2011-SCI-0048, MOJA-2013-SCI-0004, and MOJA-2016-SCI-0002. Requisite clearances and permits were obtained from the Office of Chief Wildlife Warden of Ladakh, Government of India for work in 2017.

## Supplementary Material

### Assembly of Volume Displacement Apparatus

The volume displacement (VD) apparatus was assembled using common laboratory supplies (**Diagram 1 & Figures S1A-C**); no major purchases were required for reliable and long-term operation of the apparatus. Materials included two 50 mL conical tubes, a tube rack, <3 ft of Tygon tubing (1/8 in, 1/4 in, 1/16 in), 1 one-hole rubber stopper (1.4 × 1.1 in), 1 two-hole rubber stopper (1.4 × 1.1 in), parafilm (optional), a 15 mL graduated cylinder (or a 15 or 50 mL conical tube), a mass balance (minimum accuracy of 0.01 g), a stir plate, 3 mm magnetic stir bar, and a stopwatch. Solid rubber stoppers were drilled (size 1 drill bit) to produce one or two holes (as shown **Figure S1A**).

As illustrated in **Diagram 1**, apparatus assembly entailed sequential connection of three chambers, which were respectively used for mixing (conical tube), water displacement (conical tube) and water collection (graduated cylinder or conical tube). The chambers were connected using a minimum amount of Tygon tubing (Tubes A & B), which were threaded through the rubber stoppers using needle-nose pliers. The one-hole rubber stopper was used to seal the mixing chamber (1^st^ chamber), and the two-hole rubber stopper was used to seal the displacement chamber (2^nd^ chamber). To connect the mixing and displacement chambers, Tube A (~5-6 in, ~12-15 cm) was threaded through the one- and two-hole rubber stoppers, and trimmed <2 cm from the bottom edge of each stopper. To connect the displacement and collection chambers, Tube B (~14-15 in, ~35-38 cm) was threaded through the two-hole rubber stopper, trimmed ~ 1 cm from the from the base of the mixing chamber, and inserted into the collection chamber.

**Diagram 3.**
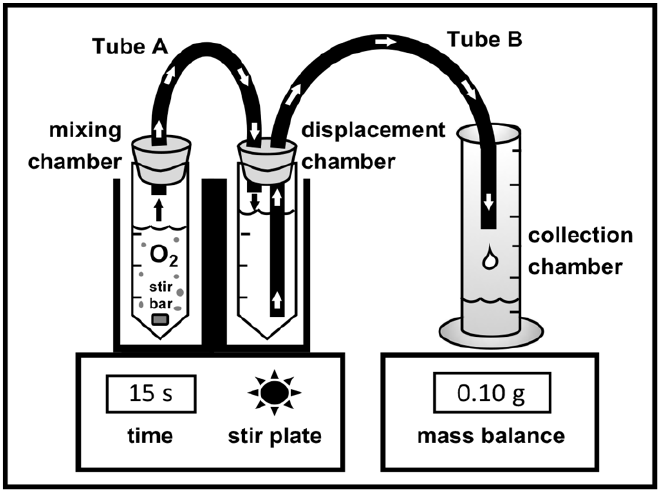
Volume displacement device with sequentially connected mixing, displacement, and collection chambers.

For the enzyme assay, the mixing and displacement chambers were inserted into a mini tube rack (as shown in **Figure S1B**) and placed on a stir plate, while the collection chamber was placed on a mass balance. Upon addition of substrate to the mixed soil suspension, the mixing chamber was immediately sealed with the one-hole rubber stopper. In turn, soil enzyme activity was monitored by following the change in mass of the collection chamber over time (alternatively, in early control experiments, changes in volume were measured using a graduated cylinder). Occasionally, the mixing chamber was wrapped in parafilm; however, no observable changes in the rates of mass or volume displacement were noted. In control experiments, priming of the connective tubing with water (with clamped control of flow into the collection chamber) yielded an ≤10% increase in rates; however, the time required per experiment commensurately increased 3-4-fold.

### Assembly of Electrochemical Apparatus

For the electrochemical analyses, a O_2_ Gas Sensor (O2-BTA) and LabQuest 2 (LABQ2) data logger were purchased from Vernier Software & Technology with respective list costs of $199 and $329. Along with these connected components, the electrochemical apparatus was assembled using a support stand (~15-24 in tall), 3-prong clamp, magnetic stir plate, 3 mm stir bar, 50 mL conical tube (mixing chamber), one and two-hole rubber stoppers, Tygon tubing, two plastic stopcocks, and parafilm.

For lower activity samples (*e.g.*, permafrost), the sensor was directly fitted into the mixing chamber (50 mL conical tube) and sealed with parafilm (**Figure S1D**). For higher activity samples (*e.g.*, BSCs and garden soils), the mixing chamber was connected to the sensor using Tygon tubing, stopcocks, and rubber stoppers. For this configuration, Tygon tubes of differing length (Tubes A-C) were prepared (Tube A: ~6-7 in, ~15-18 cm; Tube B: ~4-5 in, ~10-13 cm; Tube C: ~3-4 in; ~8-10 cm), where Tubes A and B were threaded through a two-hole rubber stopper, Tube C was threaded through a one-hole rubber stopper, and all tubes were trimmed ~ 1-3 cm from the bottom edge of the stoppers. Plastic stopcocks were attached to end of Tubes A and B (leading from the top of the stopper); while Tube C was fastened onto the opposite end of the stopcock attached to Tube A. The two-hole rubber stopper (threaded with Tubes A and B) was used to seal the mixing chamber. The one-hole rubber stopper (threaded with Tube C) was used to seal the electrochemical sensor. In this configuration, Tube A served as an exit line (for the mixing chamber) and connected to Tube C via a stopcock, Tube C connected the sensor, and tube B served as an escape line for the mixing chamber.

When conducting the kinetic assay, the sensor was secured to a support stand using a 3-prong clamp, the mixing chamber was inserted into a mini tube rack, and mixing of the suspension was accomplished using a magnetic stir plate and 3 mm stir bar. In between assay runs, the apparatus and sensor were re-equilibrated to atmospheric conditions by removing the mixing chamber from the sensor (*i.e.*, breaking the parafilm seal), or by opening the escape valve (*i.e.*, opening the stopcock on Tube B).

**Supplementary Table 1.**
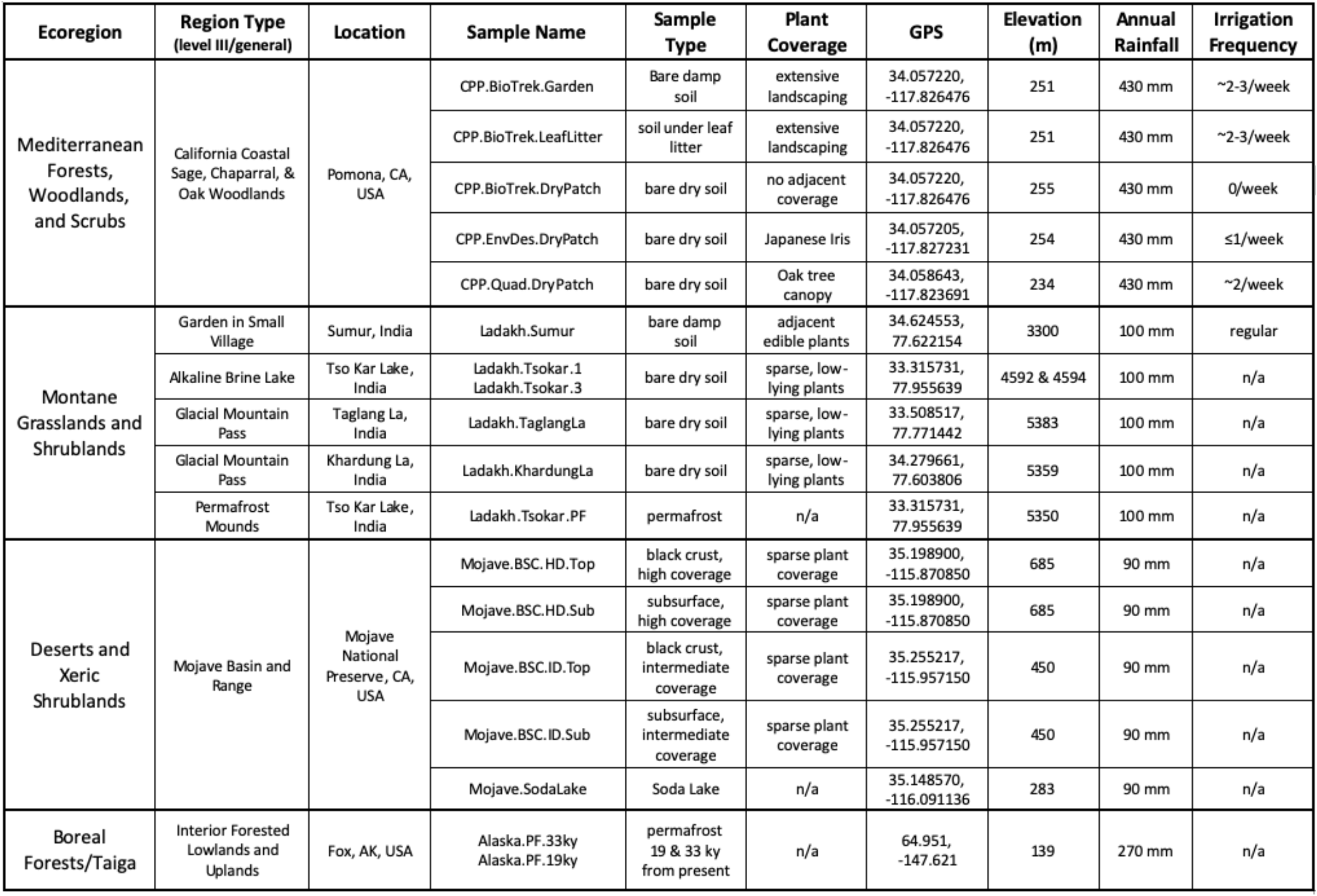
List and description of sampled soils.

**Supplementary Table 2.** List of catalase specific activities (SAs) expressed per gram dry weight (μkat gdw^−1^) and 16S rRNA copy number (fkat copy number^−1^; abbreviated as fkat 16S^−1^); where standard deviations for SAs per gdw are provided, SAs for Ladakh are expressed per gram, calculated estimates of 16S rRNA copy numbers are from a standard curve, and measured copy numbers are from Mogul et al. 2017 and Mackelprang et al 2017.

| Sample | $\mu\text{kat gdw}^{-1}$<br>(or $\text{g}^{-1}$) | stdev | Units $\text{gdw}^{-1}$<br>(or $\text{g}^{-1}$) | calc<br>16S<sup>a</sup> | measured<br>16S<sup>a</sup> | (fkat 16S<sup>-1</sup>)<sup>a</sup> |
| --- | --- | --- | --- | --- | --- | --- |
| **50 mM HEPES (pH 7.5), 330 mM non-stabilized H<sub>2</sub>O<sub>2</sub> (1% w/w)** |  |  |  |  |  |  |
| CPP.BioTrek.DryPatch (241 m) | 3.4E+00 | 2.0E-01 | 2.0E+02 | 1.4E+07 |  | 2.5E+02 |
| CPP.BioTrek.Garden (241 m) | 1.0E+01 | 7.0E-01 | 6.0E+02 | 2.3E+12 |  | 4.4E-03 |
| CPP.BioTrek.LeafLitter (241 m) | 2.2E+00 | 1.0E-01 | 1.3E+02 | 1.5E+06 |  | 1.4E+03 |
| CPP.Env.Des.DryPatch (241 m) | 6.1E+00 | 2.0E-01 | 3.7E+02 | 2.0E+09 |  | 3.1E+00 |
| CPP.Quad.DryPatch (241 m) | 4.7E+00 | 4.0E-01 | 2.8E+02 | 1.6E+08 |  | 2.9E+01 |
| Mojave.BSC.HD.Sub.DSC (284 m) | 3.3E-01 | 5.0E-02 | 2.0E+01 |  | 1.5E+07 | 2.2E+01 |
| Mojave.BSC.HD.Top.DSC (284 m) | 3.9E+00 | 2.0E-01 | 2.4E+02 |  | 5.3E+07 | 7.4E+01 |
| Mojave.BSC.HD.Top.Field (284 m) | 8.1E+00 | 5.0E-01 | 4.9E+02 |  | 5.3E+07 | 1.5E+02 |
| Mojave.BSC.ID.Sub.DSC (284 m) | 3.4E-01 | 8.0E-02 | 2.1E+01 |  | 1.1E+07 | 3.0E+01 |
| Mojave.BSC.ID.Top.DSC (284 m) | 3.5E+00 | 3.0E-01 | 2.1E+02 |  | 1.5E+07 | 2.3E+02 |
| Alaska.PF.19ky.22°C (139 m) | 1.3E+00 | 2.0E-01 | 7.5E+01 |  | 9.0E+05 | 1.4E+03 |
| Alaska.PF.33ky.22°C (139 m) | 1.4E+00 | 3.0E-01 | 8.2E+01 |  | 2.0E+05 | 6.7E+03 |
| Alaska.PF.19ky.4°C (139 m) | 9.9E-01 | 1.5E-01 | 5.9E+01 |  | 9.0E+05 | 1.1E+03 |
| Alaska.PF.33ky.4°C (139 m) | 7.0E-01 | 1.2E-01 | 4.2E+01 |  | 2.0E+05 | 3.4E+03 |
| **1x PBS, stabilized 330 mM H<sub>2</sub>O<sub>2</sub> (1% w/w)** |  |  |  |  |  |  |
| CPP.BioTrek.DryPatch (241 m) | 2.4E+00 | 2.0E-01 | 1.4E+02 | 2.4E+06 |  | 1.0E+03 |
| CPP.BioTrek.Garden (241 m) | 9.7E+00 | 6.0E-01 | 5.8E+02 | 1.4E+12 |  | 7.1E-03 |
| CPP.BioTrek.LeafLitter (241 m) | 1.9E+00 | 1.0E-01 | 1.1E+02 | 9.5E+05 |  | 2.0E+03 |
| CPP.Env.Des.DryPatch (241 m) | 2.4E+00 | 2.0E-01 | 1.4E+02 | 6.2E+07 |  | 3.9E+01 |
| CPP.Quad.DryPatch (241 m) | 4.2E+00 | 3.0E-01 | 2.5E+02 | 7.0E+07 |  | 6.0E+01 |
| Ladakh.KhardungLa (4491 m) ($\text{g}^{-1}$) | 4.5E-01 | 3.0E-02 | 2.7E+01 | 6.8E+04 | | 6.6E+03 |
| Ladakh.Sumur (3300 m) ($\text{g}^{-1}$) | 2.2E+00 | 1.1E-01 | 1.3E+02 | 2.1E+06 | | 1.1E+03 |
| Ladakh.TaglangLa (5383 m) ($\text{g}^{-1}$) | 9.1E-01 | 6.0E-02 | 5.5E+01 | 1.6E+05 | | 5.8E+03 |
| Ladakh.Tsokar.1 (4592 m) ($\text{g}^{-1}$) | 3.2E-01 | 6.0E-02 | 1.9E+01 | 5.3E+04 | | 6.0E+03 |
| Ladakh.Tsokar.3 (4594 m) ($\text{g}^{-1}$) | 3.1E-01 | 4.0E-02 | 1.9E+01 | 5.3E+04 | | 5.9E+03 |
| Ladakh.Tsokar.PF (5350 m) ($\text{g}^{-1}$) | 6.5E-02 | 3.0E-02 | 3.9E+00 | 3.4E+04 | | 1.9E+03 |
| <sup>a</sup>16S = 16S rRNA gene copy number |  |  |  |  |  |  |

**Supplementary Figure 1.**
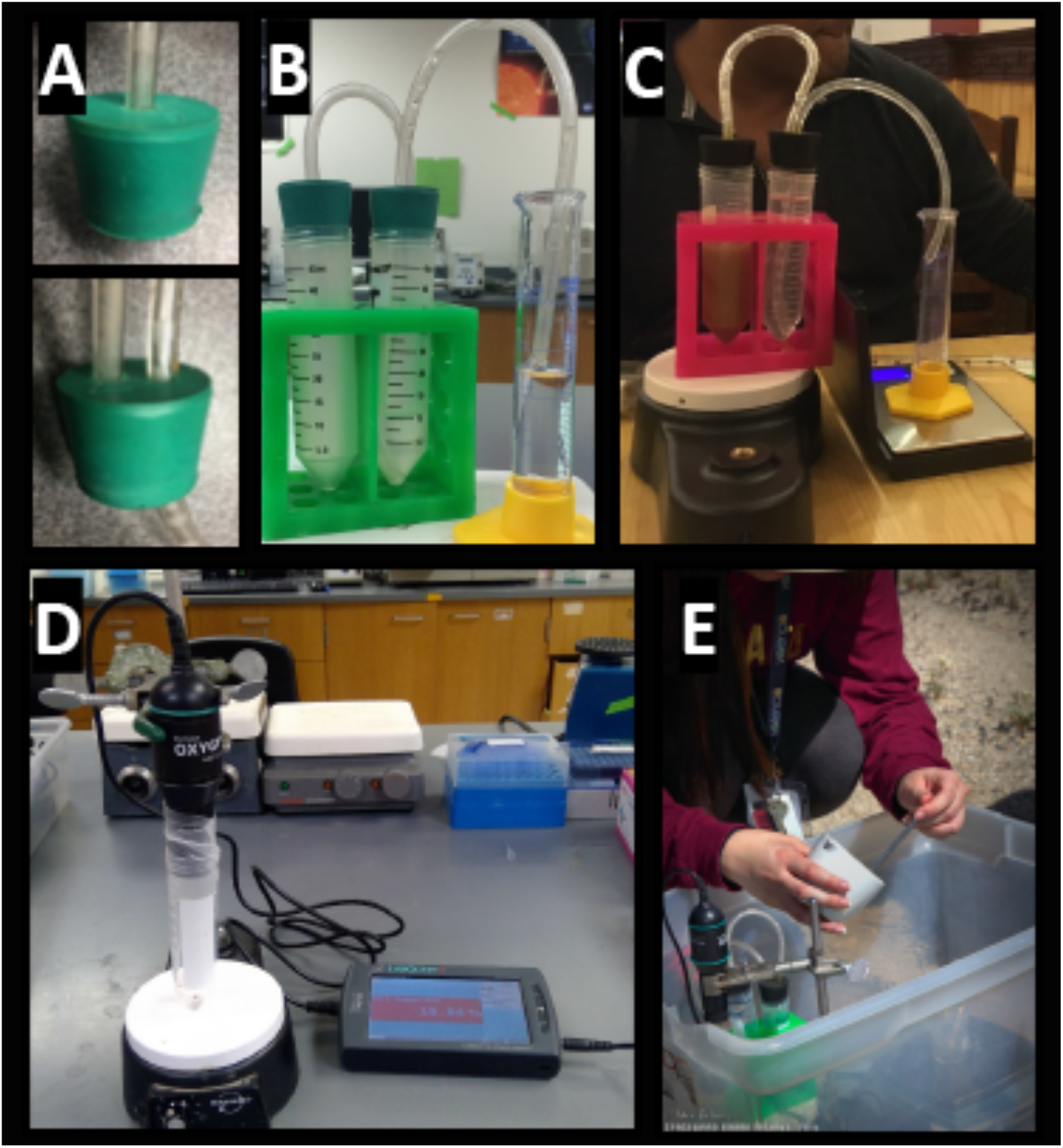
Differing configurations and components of the catalase assay devices displaying (A) one- and two-hole stoppers with threaded tubing, (B) sequential connections of the mixing, displacement, and collection chambers in the VD device (left to right), (C) battery-operated mixer and scale for VD, (D) EC device with a direct connection between the mixing chamber and sensor, and (E) example field analysis.

**Supplementary Figure 2.**
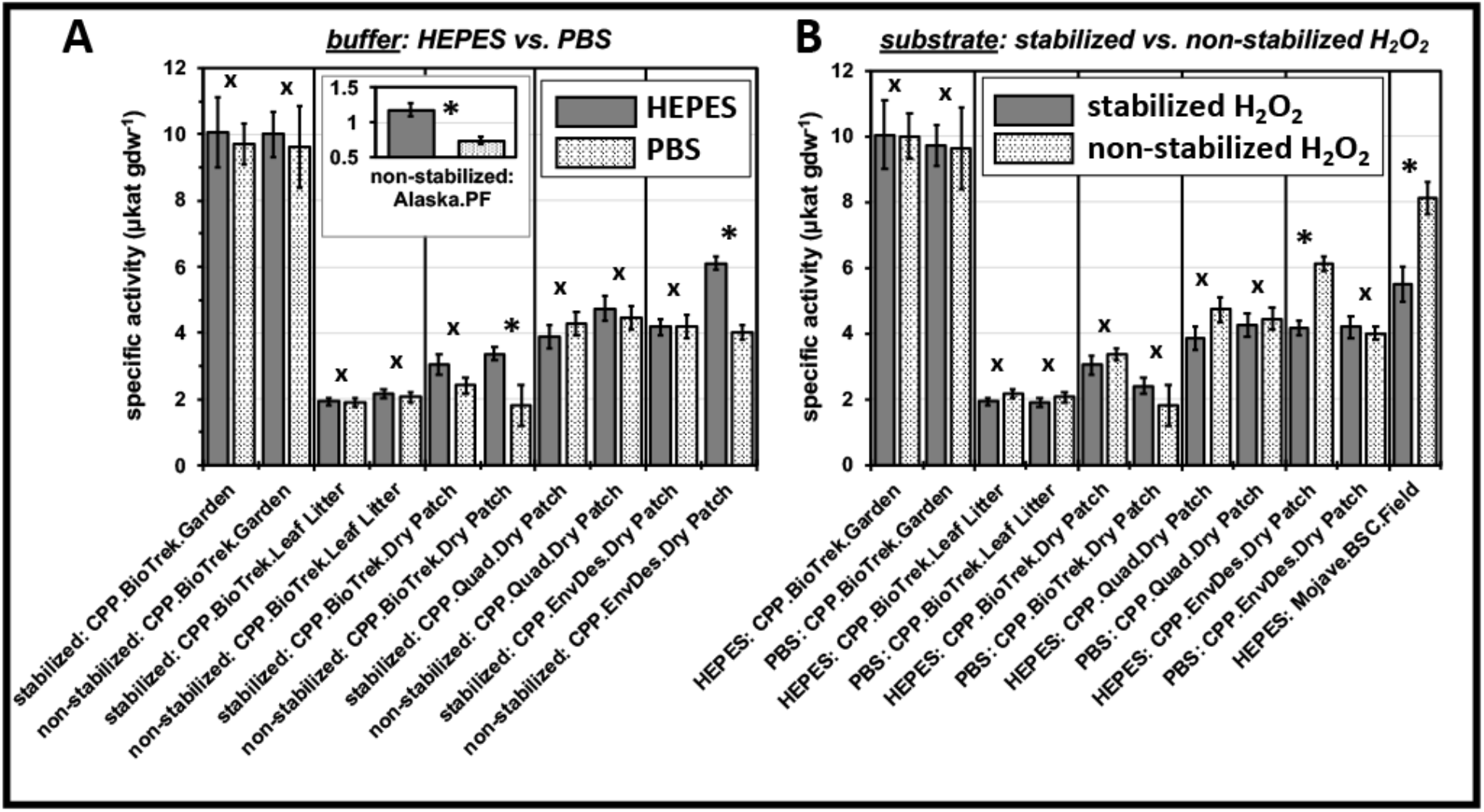
Impacts of (A) buffer type (HEPES vs. PBS) and (B) substrate formulation (stabilized vs. non-stabilized hydrogen peroxide) on the catalase specific activities, where soil samples are arranged (left to right) from highly irrigated (CPP.BioTrek.Garden) to infrequently irrigated (CPP.Env.Design.DryPatch) or arid soils (Mojave.BSC.Field), and low activity Alaskan permafrost (35k9B) is included in the inset plot; specific activities (columns) are expressed as μkat gdw^−1^, errors bars represent the propagated error (n=3), and statistical comparisons are marked (*p < 0.05; ^X^p > 0.05).

